# Variation in susceptibility among three Caribbean coral species and their algal symbionts indicates the threatened staghorn coral, *Acropora cervicornis*, is particularly susceptible to elevated nutrients and heat stress

**DOI:** 10.1101/2021.05.10.443445

**Authors:** Ana M. Palacio-Castro, Caroline E. Dennison, Stephanie M. Rosales, Andrew C. Baker

## Abstract

Coral cover is declining worldwide due to multiple interacting threats. We compared the effects of elevated nutrients and temperature on three Caribbean corals: *Acropora cervicornis, Orbicella faveolata*, and *Siderastrea siderea*. Colonies hosting different algal types were exposed to either ambient nutrients (A), elevated NH_4_ (N), or elevated NH_4_ + PO_4_ (N+P) at control temperatures (26 °C) for > 2 months, followed by a 3-week thermal challenge (31.5 °C). *A. cervicornis* hosted *Symbiodinium (S. fitti)* and was highly susceptible to the combination of elevated nutrients and temperature. During heat stress, *A. cervicornis* pre-exposed to elevated nutrients experienced 84%-100% mortality and photochemical efficiency (*F*_*v*_*/F*_*m*_) declines of 41-50%. In comparison, no mortality and lower *F*_*v*_*/F*_*m*_ declines (11-20%) occurred in *A. cervicornis* that were heat-stressed but not pre-exposed to nutrients. *O. faveolata* and *S. siderea* response to heat stress was determined by their algal symbiont community and was not affected by nutrients. *O. faveolata* predominantly hosted *Durusdinium trenchii* or *Breviolum*, but only corals hosting *Breviolum* were susceptible to heat, experiencing 100% mortality, regardless of nutrient treatment. *S. siderea* colonies predominantly hosted *Cladocopium* C1 (*C. goreaui*), *Cladocopium* C3, *D. trenchii*, or variable proportions of *Cladocopium* C1 and *D. trenchii*. This species was resilient to elevated nutrients and temperature, with no significant mortality in any of the treatments. However, during heat stress, *S. siderea* hosting *Cladocopium* C3 suffered higher reductions in *F*_*v*_*/F*_*m*_ (41-56%) compared to *S. siderea* hosting *Cladocopium* C1 and *D. trenchii* (17-26% and 10-16%, respectively). These differences in holobiont susceptibility to elevated nutrients and heat may help explain historical declines in *A. cervicornis* starting decades earlier than other Caribbean corals. Our results suggest that tackling only warming temperatures may be insufficient to ensure the continued persistence of Caribbean corals, especially *A. cervicornis*. Reducing nutrient inputs to reefs may also be necessary for these iconic coral species to survive.

## Introduction

Ocean warming is recognized as the principal threat to coral reefs in the twenty-first century (Hoegh-Guldberg et al. 2007; Hughes et al. 2017). To improve the chances of coral reef persistence, reductions in carbon emissions are imperative (IPCC 2014). However, local threats such as nutrient pollution coupled with heat stress can also play a vital role in coral survivorship. The mutualistic association between corals and dinoflagellate algae (family Symbiodiniaceae; LaJeunesse et al. 2018) is sensitive to changes in both nutrients and temperature. High nutrients can increase Symbiodiniaceae growth rates and abundance (Falkowski et al. 1993; Hoegh-Guldberg 1994; Marubini and Davies 1996), reducing the photosynthate that symbiotic algae transfer to the coral host (Hoegh-Guldberg and Smith 1989; Muscatine et al. 1989). Acute heat stress, on the other hand, can drastically reduce the abundance of algal symbionts, in a process known as coral bleaching (Weis 2008), which typically results in the mortality of the coral host if the stress is prolonged or severe (Glynn 1993).

Although elevated temperatures and nutrients can have opposite effects on symbiont densities, pre-exposure to elevated nutrients, particularly dissolved inorganic nitrogen, do not appear to mitigate bleaching, and to the contrary, can exacerbate bleaching (Wiedenmann et al. 2013; Wooldridge et al. 2017; Donovan et al. 2020) and coral mortality due to heat stress (Zaneveld et al. 2016). However, these impacts can vary by coral species (Shantz and Burkepile 2014), nitrogen source (Burkepile et al. 2019; Fernandes de Barros Marangoni et al. 2020), nutrient concentration (D’Angelo and Wiedenmann 2014), and the relative ratio of nitrogen to phosphorus (Wiedenmann et al. 2013; Morris et al. 2019). For example, pre-exposure to moderate (<5μM) levels of ammonium (Béraud et al. 2013) or phosphate (Dunn et al. 2012), but not nitrate (Fernandes de Barros Marangoni et al. 2020), can enhance coral metabolism and bleaching recovery after heat stress.

Reefs in the Caribbean have been devastated by human impacts, with relative losses of coral cover in the region since the 1970s averaging at least 50-80% (Gardner et al. 2003; Jackson et al. 2014). Early coral declines were attributed to decreasing water quality (Smith et al. 1981; Tomascik and Sander 1987; Lapointe and Clark 1992), overfishing (Hughes 1994; Jackson et al. 2001), and disease (Aronson and Precht 2001; Schutte et al. 2010). However, the role of thermal stress-induced coral bleaching (Williams et al. 1987; Aronson et al. 2002; McWilliams et al. 2005; Lough et al. 2018) and its interaction with local stressors (Zaneveld et al. 2016; Hughes et al. 2017; Wang et al. 2018; Lapointe et al. 2019) has been increasingly recognized since the late 1980s (Williams et al. 1987; Glynn 1993).

Importantly, cover loss has not been homogeneous among coral species, resulting in changes in species dominance and reef structure over time (Aronson et al. 2004). Caribbean acroporids (*Acropora cervicornis* and *A. palmata*), the only fast-growing taxa with branching morphologies in the region, were the earliest taxa to show significant declines and were listed as threatened by the US Endangered Species Act in 2006. About 90% of acroporids were lost from the 1970s to the 1990s due to disease and bleaching (Aronson and Precht 2001; Aronson et al. 2002), but recent studies suggest that their decline may have started 2-3 decades earlier (Cramer et al. 2020). Similarly, significant reductions in *Orbicella* spp. were documented in multiple locations during the 1990s and 2000s (Bruckner and Hill 2009), and as a result, *O. annularis, O. faveolata*, and *O. franksi* were listed as threatened in 2014. Other coral taxa, such as *Agaricia, Porites*, and *Siderastrea* have been more resistant to stress, and have increased their relative abundance on some reefs (Goreau 1992; Aronson et al. 2004; Pandolfi and Jackson 2006). However, unlike *Acropora* and *Orbicella*, these corals are not primary builders of reef frameworks, and consequently do not provide the same ecosystem services of more structurally complex coral species (Alvarez-Filip et al. 2013; De Bakker et al. 2016).

In this study, we examined the effects of pre-exposure to elevated nutrients (NH_4_ and NH_4_ + PO_4_ for >2 months at 26 °C), followed by heat stress (31.5 °C for 3-weeks) on three Caribbean corals. Two of the species (*A. cervicornis* and *O. faveolata*) are listed as threatened under the Endangered Species Act, and one (*S. siderea*) is a hardy species that, although slow-growing, is nevertheless still relatively common on Caribbean reefs. We aimed to compare the effects of these combined stressors on coral survivorship, and associated algal symbiont communities (community composition, abundance, and function). Our goal was to better understand the differential impacts of nutrient pollution and heat stress on different coral holobionts, and to gauge how these stressors might continue to shape Caribbean reefs over the next decades if they are not effectively managed.

## Methods

### Coral collection

Replicate fragments of *A. cervicornis, O. faveolata*, and *S. siderea* were collected in the summer of 2017. *A. cervicornis* (N=6 genets) were obtained as single branches (n=30-31 per genet, ∼ 4 cm long) from coral nurseries operated by the University of Miami and Mote Marine Laboratory. Whole colonies of *O. faveolata* (N=4) and *S. siderea* (N=7) were collected from Emerald Reef (off Key Biscayne, FL, 25.673°N, 80.098°W) under a Special Activity License (SAL-17-1182-SRP) and were subdivided in the laboratory using a drill press to obtain replicate cores (n=18-24 per colony, ∼ 2 -2.5 cm diameter). All fragments were maintained in indoor tanks at ∼ 26°C for at least two months for recovery.

### Experimental conditions

On day one of the experiment, we collected biopsy samples and baseline photochemical efficiency data from each coral fragment, and haphazardly assigned them to a nutrient treatment in one of two replicate tanks. Corals in the ambient treatment (A) were exposed to nutrient levels typical for Biscayne Bay, FL. Corals in elevated NH_4_ (N) were exposed to a 10 μM increase in ammonium, and corals in elevated NH_4_ + PO_4_ (N+P) were exposed to 10 μM increase in ammonium, plus a 1 μM increase in PO_4_. Each treatment was replicated in two independent 38-L glass aquaria immersed in two fiberglass tanks, which acted as water baths to maintain target temperatures. This setup resulted in 2-10 fragments per colony exposed to each nutrient treatment (1-5 fragments per colony in each replicate aquarium (Table S1). Three times per week, all corals were transferred to a separate raceway and where they were fed for 1 h with Zeigler® AP 110 dry larval diet. Before returning the fragments to their experimental aquaria, fragments and treatments were rotated in the tanks to randomize the effect of variation in light intensity across the tanks (which ranged from 321 - 420 μmol PAR m-2 s-1). More detailed methods are described in ESM1.

For ∼2.5 months (78 d) the corals were maintained at control temperature (26.1 ± 0.4 °C; mean ± SD). On day 79, the temperature was incrementally increased over 12 d (“ramp-up” phase, days 79-90) until the target heat stress temperature of 31.5 °C was reached. During the next three weeks (days 91-113) corals remained in the “heat stress” phase (31.5 ± 0.8 °C). Nutrient addition ended on day 91 due to the onset of mortality in *A. cervicornis* exposed to N and N+P. Therefore, during heat stress, the corals assigned to N and N+P treatments were maintained under ambient nutrients (Fig. 1).

**Figure 1:**
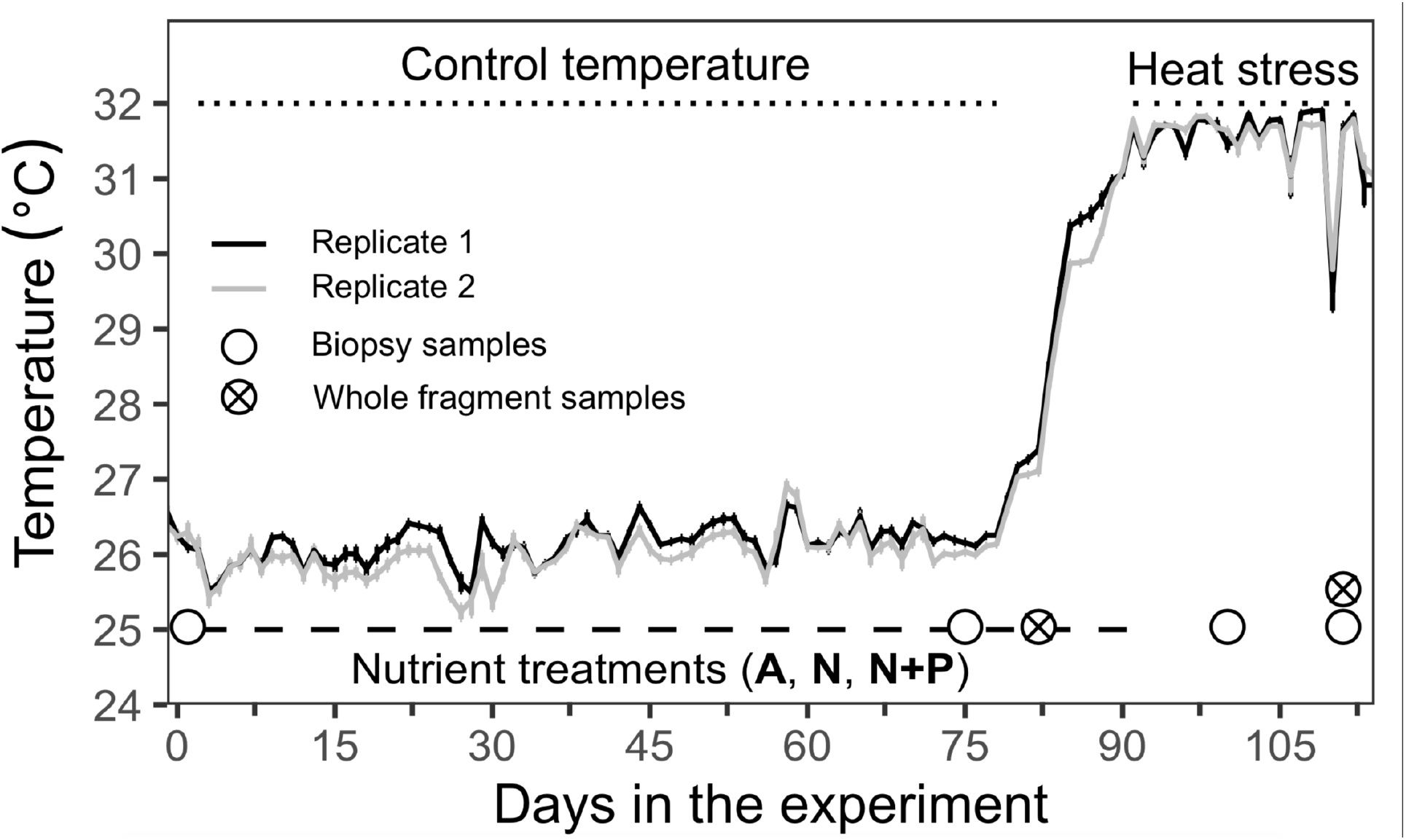
Experimental conditions during the control, ramp-up, and heat stress phases. Solid lines (black and gray) show the temperature in two replicate tanks (mean ± 95% CI), each one holding one aquarium at ambient nutrients (A), elevated ammonium (N), and elevated ammonium and phosphate (N+P). The bottom dashed line denotes the period of nutrient addition. Circles indicate the days when samples were collected.

### Coral survivorship

Mortality of fragments was recorded daily. Survival probabilities were calculated for each coral species using the Kaplan-Meier estimate (Kaplan and Meier 1958), which uses the state of each fragment at a given time (alive, dead, or unknown) to calculate the proportion of surviving individuals in the treatments. Log-rank tests were used to compare the survivorship curves for corals in A, N, and N+P during control and heat stress temperatures. For *O. faveolata* and *S. siderea*, we also tested if the dominant Symbiodiniaceae type affected survivorship. When collecting chlorophyll-*a* and symbiont density (see sections below), selected fragments were sacrificed and permanently removed from the tanks. These individuals were included in the model until their removal, but they were not considered as part of the sample groups after removal These removal events are treated as “censored” or unknown events in the model. Statistical analyses were performed with the survival 2.38 (Therneau 2015) and survminer 0.4.6 (Kassambara et al. 2019) packages for R.

### *Algal symbiont community function* (F_v_/F_m_)

The photochemical efficiency of photosystem II (*F*_*v*_ */F*_*m*_) was used as a proxy for algal symbiont community function and was monitored with a MAXI Imaging Pulse Amplitude Modulated (IPAM) fluorometer (Walz, Effeltrich, Germany). A single IPAM measurement per fragment was taken every two weeks during the control temperature phase, and twice a week during heat stress. All corals were dark-adapted for 30 minutes before taking measurements.

### *Algal symbiont areal densities, and chlorophyll-*a *concentrations*

A subset of fragments was removed from the tanks before (day 78) and after heat stress (day 113) to estimate symbiont areal densities (no. of symbiont cells cm^-2^) and chlorophyll-*a* content (μg Chl-*a* cm^-2^) (Fig. 1). Two fragments per nutrient treatment, colony, and temperature phase were haphazardly chosen and preserved at -80°C until the tissue was processed for each analysis. Briefly, the tissue from each fragment was removed with an airbrush and DNA buffer (10 mM Tris, 1 mM EDTA). The resulting slurry was collected, homogenized, and divided between two aliquots. One aliquot was filtered onto a Whatman glass microfiber GF/F filter and then transferred to tubes containing methanol. These samples were stored at -20°C for 24 h to extract chlorophyll-a and then were measured with a TD-700 Turner fluorometer. The second aliquot was preserved and stained with 50µL of Lugol’s iodine solution. This sample was used to count the symbiont cells, using an inverted microscope and a hemocytometer. Detailed methods are presented in the ESM1. For *A. cervicornis*, fewer fragments were sampled after heat stress due to previous coral mortality in corals exposed to N and N+P (Table S2).

### Symbiodiniaceae community composition

The proportion of each Symbiodiniaceae genera in the coral fragments was assessed before nutrient addition (day 1), at the end of the control phase (day 75), and during the heat stress phase (days 100 and 111) using real-time PCR (qPCR) assays. Biopsy samples (1-3 polyps per fragment) were preserved by incubating the tissue for 90 min at 65°C in 500 μL of a solution of 1% SDS and DNA buffer (Rowan and Powers 1991) and genomic DNA was extracted from the SDS lysates using standard procedures (Baker and Cunning 2016).

TaqMan master mix (Thermo Fisher Scientific) assays were used to amplify the actin gene in *Breviolum* (Cunning et al. 2015), *Cladocopium*, and *Durusdinium* (Cunning and Baker 2013). SYBR-Green master mix (Thermo Fisher Scientific) assays were used to quantify *Symbiodinium*, using the A_actin primers described in (Winter 2017). All coral hosts were quantified with SYBR-Green assay and primers that target species-specific single-copy loci. Calmodulin-gene (CaM) primers (CaM_forward: 5’-GCCCTAATTTCTGATCGATTCAA-3’, CaM_reverse: 5’-GCAGACAGAAGGGCCACT-3’) were developed and validated for *A. cervicornis* (ESM1). OfavscF1 and OfavscR1 (Cunning et al. 2015) were used for *O. faveolata*, and Pax-C_F and Pax-C_R (Cunning 2013) were used for *S. siderea*. Reagents and qPCR conditions for each assay are described in ESM1. The qPCR data (C_T_ values) were used to estimate the symbiont to host cell ratio (S/H ratio; Mieog et al. 2009; Cunning and Baker 2013) using the StepOne package for R (Cunning 2018). For corals hosting multiple Symbiodiniaceae genera, the total symbiont to host cell ratio (S/H) was calculated as the addition of all genera cell ratios (S/H = *Symbiodinium*/Host + *Breviolum*/Host + *Cladocopium*/Host + *Durusdinium*/Host), and the proportion of the symbiont community composed by thermotolerant *D. trenchii* was calculated as (*D. trenchii*/Host)/(S/H).

Additionally, a subset of DNA samples from *S. siderea* colonies that hosted *Cladocopium* were used to identify the specific *Cladocopium* lineage based on the nuclear ribosomal internal transcribed spacer region 2 (ITS2). ITS2 sequences from excised dominant bands in DGGE gels (LaJeunesse and Trench 2000; LaJeunesse 2002) were aligned and compared with published sequences available in GenBank. *S. siderea* and *O. faveolata* colonies predominantly hosting *Durusdinium* were not verified with ITS2 but were assumed to harbor *D. trenchii*, the only *Durusdinium* species identified to date from the Caribbean (Pettay et al. 2015). Similarly, *A. cervicornis* colonies hosting *Symbiodinium* were assumed to be *Symbiodinium* A3, also known as *S. fitti*, the only member of this genus known to associate with *A. cervicornis* (LaJeunesse 2002; Thornhill et al. 2006).

### Statistical analyses

The effects of elevated nutrients during control temperatures, and pre-exposure to nutrients during heat stress were analyzed using mixed-effects models with the lme4 1.1-17 package (Bates et al. 2015) for R 3.4.3 (R Core Team 2018). Changes in *F*_*v*_ */F*_*m*_, symbiont areal densities, and chlorophyll-*a* concentrations were individually tested for each coral species with models that included *nutrient* treatment, dominant *symbiont* type, and *days* in the experiment as interacting fixed factors. Coral *genotype* and *replicate tank* were included as random effects in all models. Additionally, *fragment* was included as a random factor in the *F*_*v*_ */F*_*m*_ models. Pairwise comparisons between significant effects were performed with the package emmeans 1.1.3 (Lenth 2018). An alpha value of 0.05 was pre-set as the threshold for the Tukey’s HSD contrasts. Models and pairwise tables are presented in ESM3. All figures show the mean values for the variables (± 95% CI) and were created with ggplot2 (Wickham 2016).

## Results

### Molecular identification of algal symbionts

All *A. cervicornis* genets hosted the genus *Symbiodinium*, which was assumed to be *S. fitti* (see Methods). No other Symbiodiniaceae genus was detected in *A. cervicornis*. In *O. faveolata*, three colonies predominantly hosted *Durusdinium* (*Durusdinium* proportion > 0.99), and one predominantly hosted *Breviolum* (*Breviolum* proportion > 0.99). However, three fragments from this colony were found to have relatively even amounts of *Durusdinium* and *Breviolum* (ESM2-Fig. S1), suggesting spatial variability in symbiont community structure within this colony (Rowan et al. 1997). Nutrients did not change the relative abundance of the different symbiont genera in *O. faveolata*. However, during heat stress (day 100) all but one fragment that predominantly hosted *Breviolum* and survived became dominated by *Durusdinium* (ESM2-Fig. S1). *Breviolum* or *Durusdinium* subtypes were not further identified using ITS-2, but colonies hosting *Durusdinium* were assumed to be *D. trenchii* (Pettay et al. 2015). In *S. siderea*, three colonies predominantly hosted *D. trenchii*, two colonies *Cladocopium* C3, one colony *Cladocopium* C1 (*C. goreaui*, LaJeunesse et al. 2018), and one colony had variable proportions of *Cladocopium* C1 and *D. trenchii*. Nutrients did not affect the relative abundance of the different symbiont genera hosted by *S. siderea*. However, during heat stress, some cores initially dominated by *Cladocopium* C3 became dominated by *D. trenchii* (ESM2-Fig. S2).

### Coral survivorship

Elevated nutrients reduced survivorship in *A. cervicornis*, but not in *O. faveolata* or *S. siderea*. However, *A. cervicornis* mortality was more pronounced under elevated nutrients and heat stress combined (Fig. 2). In *A. cervicornis* there was no mortality during the first two months at control temperature (∼26 °C) in any of the nutrient treatments. However, by days 65-78, *A. cervicornis* exposed to N and N+P experienced rapid tissue loss (ESM 2-Fig. S3), which reduced survivorship probabilities by 10% in N (p<0.05) and 3% in N+P (p>0.05). Mortality was exacerbated during ramp-up and heat phases but only affected fragments that had been pre-exposed to N and N+P. After one day at 31.5°C, survivorship probabilities were 30% and 12% lower in *A. cervicornis* pre-exposed to N and N+P, respectively, and were 100% and 84% lower after 3 weeks (Fig. 2; Log-rank p<0.0001). Although mortality was higher in N than in N+P, survivorship probabilities were not significantly different between these treatments (Log-rank p=0.097). None of the *A. cervicornis* fragments maintained in ambient nutrients (A) died during the control or heat stress phases.

**Figure 2:**
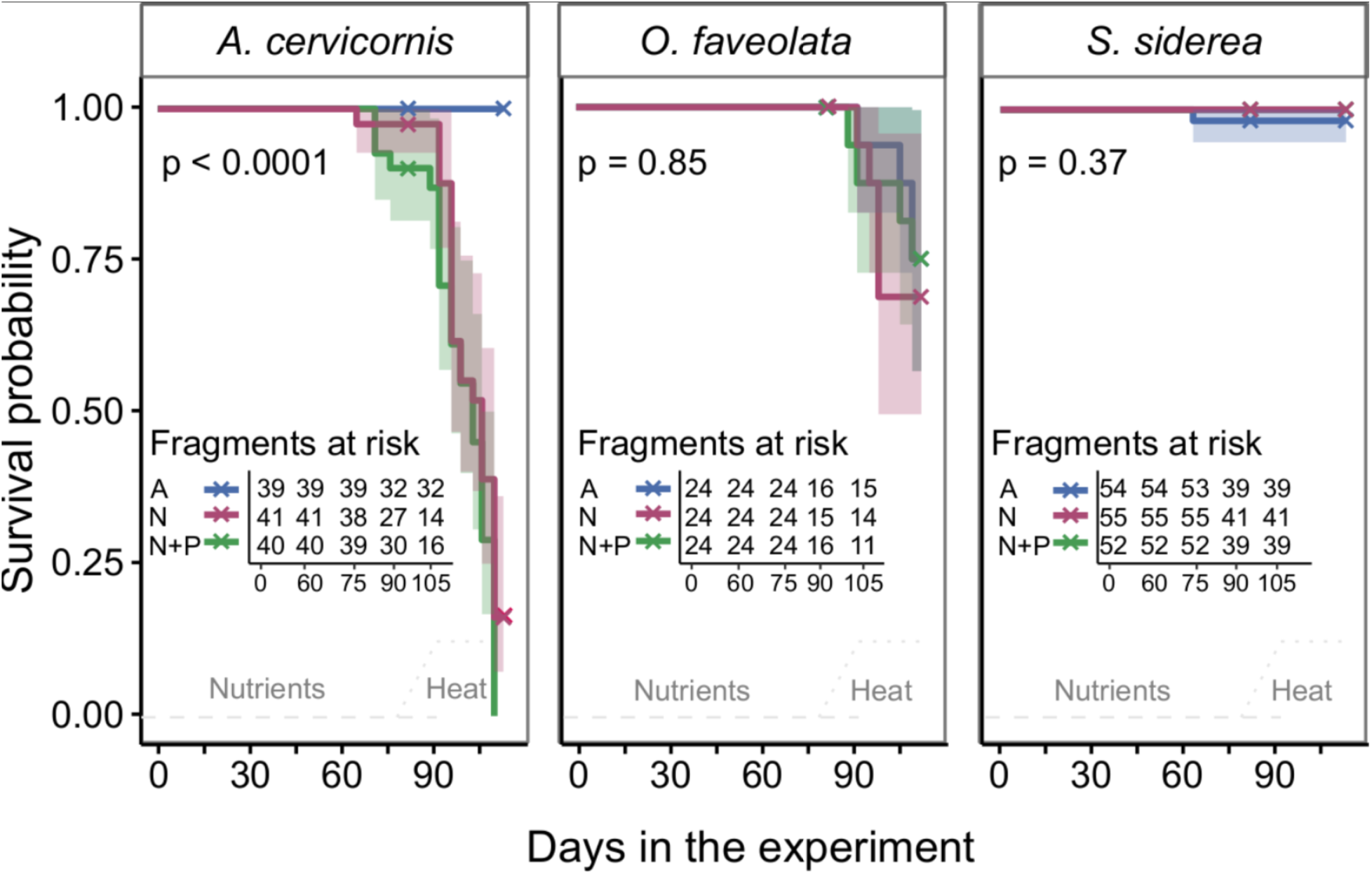
Survivorship probabilities of *Acropora cervicornis, Orbicella faveolata, and Siderastrea siderea*. Each panel represents a single species under each nutrient treatment (A, N, and N+P) during the control temperature (days 2-78), ramp-up (days 79-90), and heat stress phases (days 91-113). Fragments at risk tables show each nutrient treatment (y-axis), the days in the experiment (x-axis), and the respective number of fragments that remained in the experiment on any specific day (initial number of fragments minus fragments that died or were removed to collect whole-tissue samples). The “x” symbols represent “censored” events (days when fragments were removed to obtain samples). The bottom dashed line denotes the period of nutrient addition and the dotted lines denote the ramped-up and heat stress periods.

Heat stress (alone or combined with elevated nutrients) reduced survivorship in *O. faveolata*, but this effect depended on the symbiont type hosted. During heat stress, the colony hosting *Breviolum* experienced significant mortality (ESM2-Fig. S4; Log-rank p<0.0001) independent of nutrient treatment (Fig. 2; Log-rank p=0.85). Survival probabilities in this colony were ∼33% lower after one day under heat stress, and 100% lower after 3 weeks (ESM2-Fig. S4). *S. siderea* did not experience significant mortality during the control or heat stress phases in any of the nutrient treatments (Fig. 2; Log-rank p=0.37).

### *Symbiont community function* (F_v_ /F_m_)

During the first 1-2 months at control temperatures, elevated nutrients (N and N+P) increased photochemical efficiency (*F*_*v*_*/F*_*m*_) in *A. cervicornis + S. fitti* (+5-10%), *O. faveolata* + *D. trenchii* (+8-13%), and *S. siderea + D. trenchii* (+8-13%) compared to ambient nutrients (A) (Fig. 3; Tukey’s HSD p<0.05). The opposite effect was observed for *S. siderea + Cladocopium* (C1 or C3), which, by the end of the control phase (days 65-71), had ∼ 8-15% lower *F*_*v*_ */F*_*m*_ in fragments maintained in N compared to A (Fig. 3; Tukey’s HSD p<0.05).

**Figure 3:**
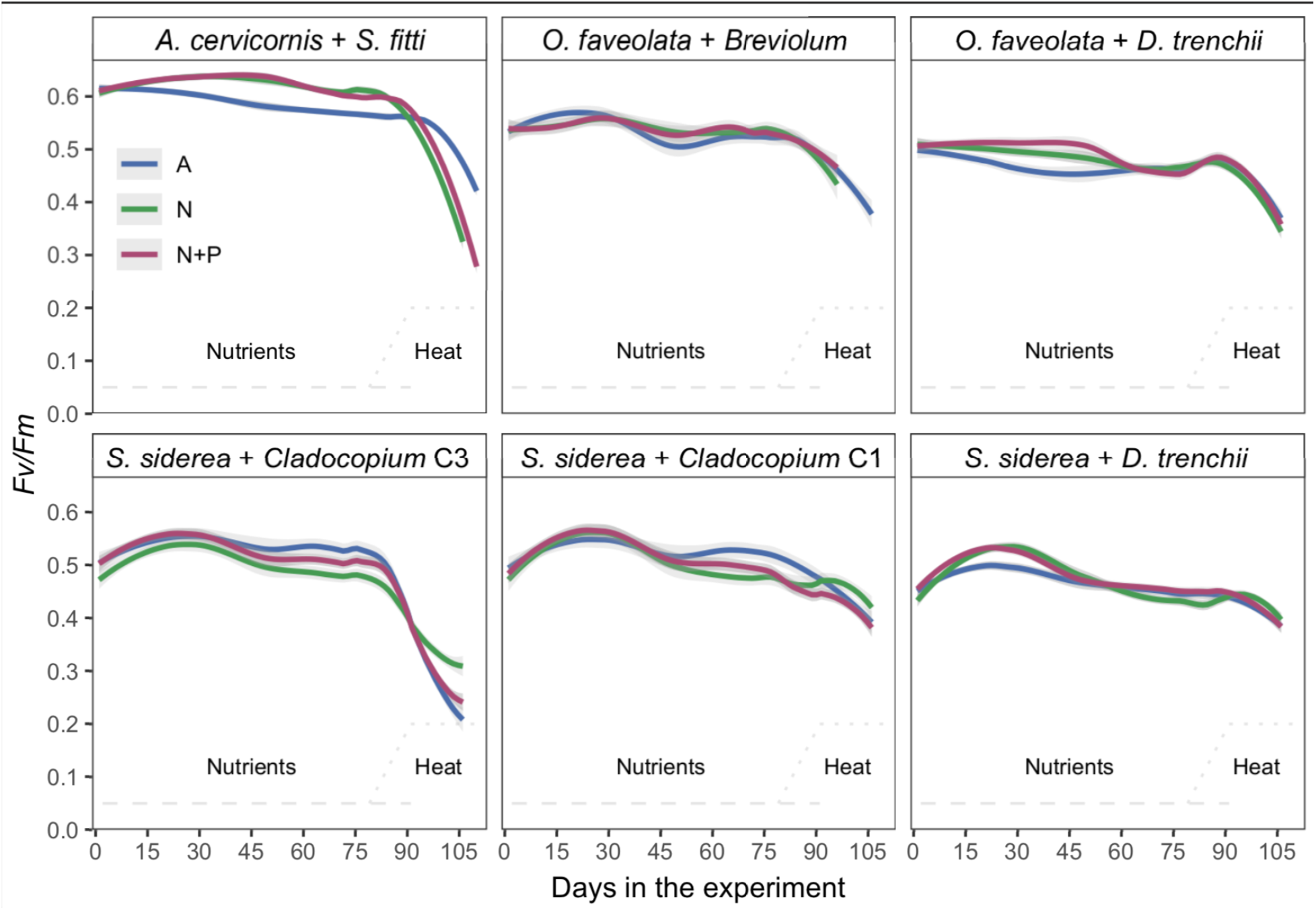
Photochemical efficiency of *A. cervicornis, O. faveolata, and S. siderea* by symbiont community. Each panel represents the photochemical efficiency (mean *F*_*v*_ */F*_*m*_ ± 95% CI) of a single coral species and dominant algal symbiont under each nutrient treatment (A, N, and N+P) during control, ramp-up, and heat stress phases. The data were fitted to a smooth line trend (span = 0.5) to highlight the overall trends. The bottom dashed line denotes the period of nutrient addition and the dotted lines denote the ramp-up and heat stress periods.

Heat stress reduced *F*_*v*_ */F*_*m*_ in all coral holobionts, but the magnitude varied by coral species, dominant algal symbiont type, and pre-exposure to nutrients (Fig. 3). During heat stress, *S. siderea* + *Cladocopium* C3 showed the earliest and largest declines in *F*_*v*_ */F*_*m*_ overall. After one day in heat, *Cladocopium* C3 fragments had experienced 18-23% reduction in *F*_*v*_ */F*_*m*_ with respect to controls (Tukey’s HSD p<0.05); and after one week, declines reached more than 40% (−41% in N, -54% in A, and -56% in N+P; Tukey’s HSD p<0.05). Compared to *Cladocopium* C3, fragments that hosted *Cladocopium* C1 and *D. trenchii* experienced smaller reductions in *F*_*v*_*/F*_*m*_ during heat stress. *Cladocopium* C1 corals experienced a maximum *F*_*v*_ */F*_*m*_ decline of 17-26% (Tukey’s HSD p<0.05), while cores hosting *D. trenchii* had a maximum decline of 10-16% (Tukey’s HSD p>0.05).

*A. cervicornis + S. fitti* had the second largest reduction in *F*_*v*_ */F*_*m*_. However, in these corals, the effect of heat stress was exacerbated by pre-exposure to elevated nutrients. Corals exposed to A, N, and N+P experienced 5%, 32%, and 18% reductions in *F*_*v*_*/F*_*m*_, respectively, after one week in heat compared to control temperatures (Tukey’s HSD p<0.05). After two weeks in heat stress, *F*_*v*_ */F*_*m*_ declined by 11%, 49%, and 41% (Tukey’s HSD p<0.05; Fig. 3).

In *O. faveolata + D. trenchii, F*_*v*_ */F*_*m*_ declines with respect to values at control temperature were significant after two weeks in heat stress (17-25%; Tukey’s HSD p<0.05), and pre-exposure to nutrients did not have a significant effect. There were no fragments hosting *Breviolum* at this time since they had either died or become dominated by *D. trenchii*.

### Chlorophyll-a concentrations

During the control phase (day 78), *A. cervicornis* chlorophyll-*a* concentrations in ambient (A) were 3.6 ± 0.4 μg cm^-2^ (mean ± SE). Exposure to nutrients increased these values by 43% in N (Tukey’s HSD p<0.05), and 39% in N+P (Tukey’s HSD p=0.05). After three weeks in heat stress (day 113), chlorophyll-*a* declined by 62% in A, and 91% in N+P with respect to their pre-heat stress values (Tukey’s HSD p<0.05; Fig. 4). Similarly, at control temperatures, *O. faveolata* chlorophyll-*a* concentration was 49% higher in N and 55% in N+P with respect to A (3.8 ± 0.6; Tukey’s HSD p<0.05). Heat significantly reduced these values by 40% in A, 73% in N, and 53% in N+P with respect to their pre-heat stress values (Tukey’s HSD p<0.05; Fig. 4).

**Figure 4:**
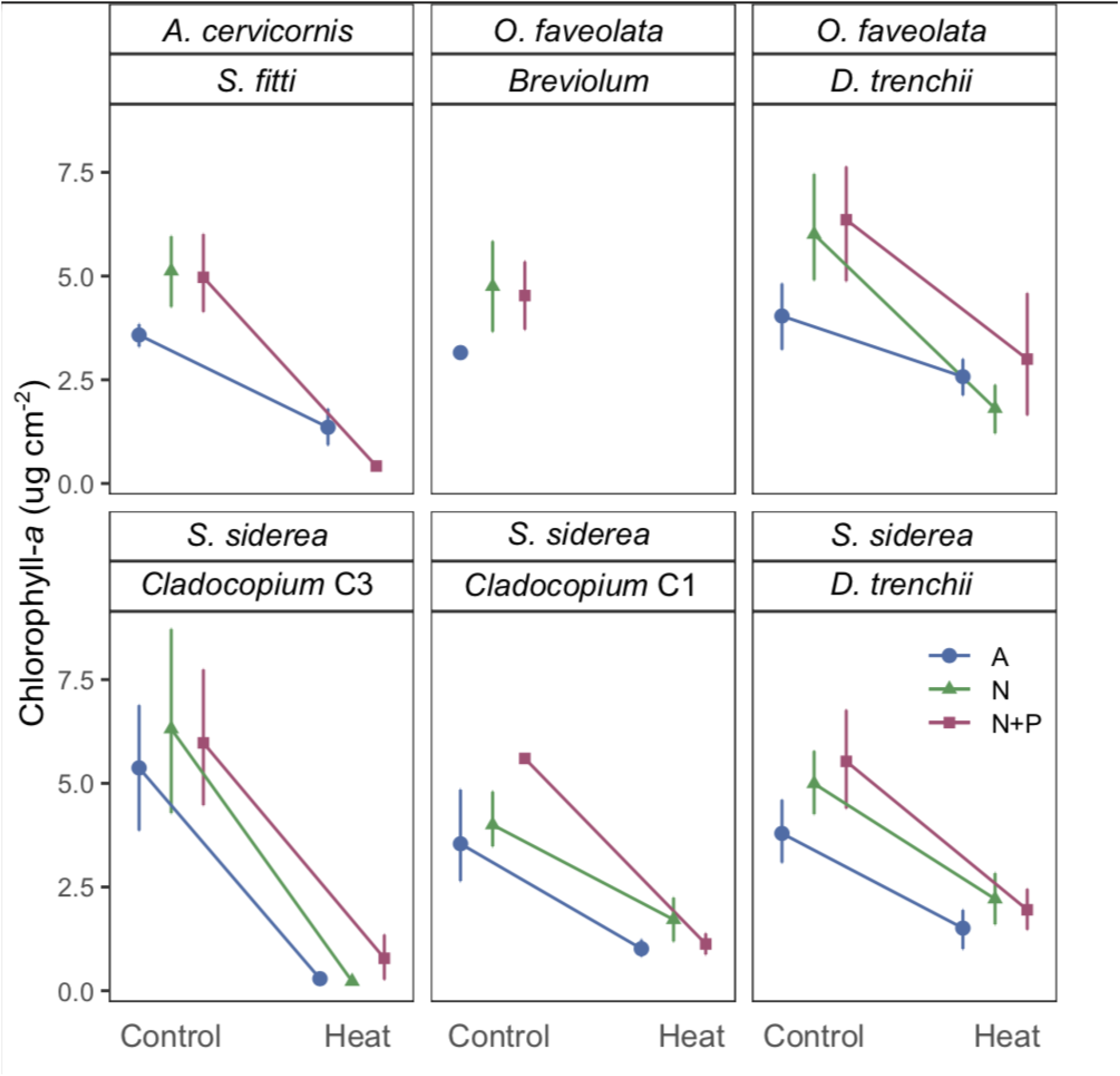
Chlorophyll-*a* concentration in *A. cervicornis, O. faveolata, and S. siderea* by symbiont community. Each panel represents the chlorophyll-*a* concentration (mean μg cm^-2^ ± 95% CI) of a single coral species and dominant algal symbiont under each nutrient treatment (A, N, and N+P) at control temperature (day 78) and heat stress (day 113).

In *S. siderea*, chlorophyll-*a* concentration varied with symbiont identity and heat stress interactions (Fig 4). At control temperature, *Cladocopium* C3 fragments had higher chlorophyll-*a* concentration (6.8 ± 0.7) than *Cladocopium* C1 (3.9 ± 0.7) and *D. trenchii* fragments (4.4 ± 0.5) (Tukey’s HSD p<0.05). Heat stress reduced chlorophyll-*a* concentration in all treatments and symbionts (Tukey’s HSD p<0.05; Fig. 4). Although chlorophyll-*a* loss was higher in Cladocopium C3 and C1 (−73.6% and -73.3%, respectively) than in *D. trenchii* (−65.5%) the final (day 113) concentrations were similar among all symbionts.

### Areal symbiont densities

Heat was the only factor that significantly affected symbiont areal densities (mean cells cm^-2^ ± SE) across all coral-algal combinations (Fig. 5). Under control temperature, *A. cervicornis* in ambient (A) hosted 4.1± 0.4 x 10^6^ cells cm^-2^. Although symbiont densities were 18.1% and 29.7% higher in *A. cervicornis* exposed to N and N+P, respectively, these differences were not significant (p>0.05 Tukey’s HSD; Fig. 5). In *A. cervicornis*, heat reduced symbiont densities by 73.9% in A (Tukey’s HSD p<0.05). Fragments in N and N+P could not be assessed due to mortality in these treatments.

**Figure 5:**
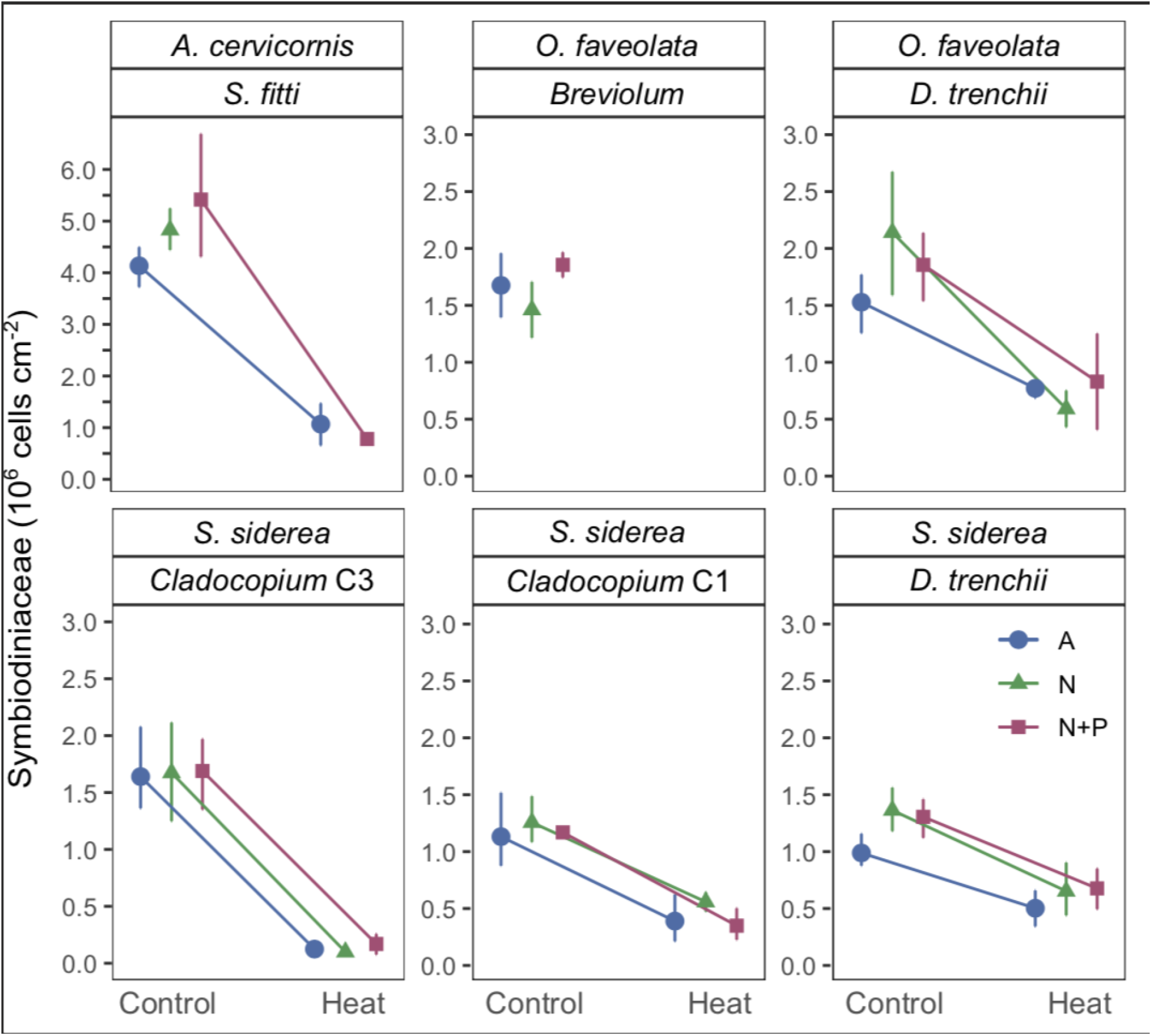
Symbiodiniaceae areal densities in *A. cervicornis, O. faveolata, and S. siderea* by symbiont community. Each panel represents the Symbiodiniaceae areal densities (mean Symbiodiniaceae cells cm^-2^ ± 95% CI) of a single coral species and their dominant algal symbionts under each nutrient treatment (A, N, and N+P) at control temperature (day 78) and heat stress (day 113).

At control temperatures, *O. faveolata* symbiont densities were similar among nutrient treatments and symbiont types (1.6 ± 0.2 x 10^6^). Heat significantly reduced symbiont densities in all nutrient treatments compared to their values at control temperature (−50% in A, -70% in N, and -59% in N+P; Tukey’s HSD p<0.05, Fig. 5).

*S. siderea* symbiont density was influenced by dominant symbiont, but not by nutrient treatment. At control temperatures, fragments hosting *Cladocopium* C3 (1.32 ± 0.1 x 10^6^) had 23% higher densities than cores hosting *Cladocopium* C1 and *D. trenchii* (Tukey’s HSD p<0.05). Heat stress reduced areal densities in all *S. siderea* symbiont combinations (Tukey’s HSD p<0.05), but symbiont loss was higher in *Cladocopium* C3 (−68%) than in *Cladocopium* C1 (−44%) and *D. trenchii* (−32%).

## Discussion

The goal of this study was to compare the response of three Caribbean corals and their associated algal symbionts to (1) elevated nutrients (N and N+P) at control temperature (∼26 °C), (2) elevated temperature (∼31.5 °C) at ambient nutrients (A), and (3) the combined effects of elevated nutrients and temperature. Of note, our experiment was designed to test the interactions between heat and nutrients, but we ceased the addition of nutrients during heat stress, due to the rapid mortality observed in *A. cervicornis* exposed to elevated N and N+P (Figs. 1-2). Despite this limitation, we were able to document different responses among multiple coral-algal combinations to elevated nutrients, heat stress, and pre-exposure to elevated nutrients followed by heat stress, which may help explain differential changes in coral cover among Caribbean coral species.

### A. cervicornis *is highly susceptible to elevated nutrients*

In our study, *A. cervicornis* was highly susceptible to elevated nutrients compared to *O. faveolata* and *S. siderea*. For example, *A. cervicornis* experienced mortality in fragments exposed to N and N+P during the control, ramp-up, and heat stress phases (Fig. 2), while mortality in *O. faveolata* only occurred during heat stress and was dependent on symbiont identity, but not previous exposure to high nutrients (ESM2-Figs. S1, S4). Although nutrients are known to exacerbate the severity of disease in corals (Bruno et al. 2003; Vega Thurber et al. 2013), the mortality of *A. cervicornis* in our study was likely caused by the disruption of algal symbiosis under the elevated nutrients, rather than other pathologies (e.g., white band or other infectious diseases). In this study, tissue loss began in different areas of the fragments (not only the bases) and tissue sloughing quickly advanced in a variety of directions, rather than as a clear band moving up from the base (ESM2-Fig. S3). Additionally, all *A. cervicornis* fragments, regardless of nutrient treatment, were fed 2-3 times a week in the same feeding chamber for ∼1 h, but mortality did not spread from corals in the N and N+P to corals in the A treatments, even during heat stress, when corals are likely to become immunocompromised (Muller et al. 2018). This suggests that the cause of death was not infectious or transmissible through the water column.

Although in our experiment pre-exposure to elevated nutrients exacerbated the effects of heat in *A. cervicornis*, it is possible that the interaction among these two stressors would have a different outcome if heat stress was followed by elevated nutrients. In this case, nutrient availability could boost bleaching recovery by providing the algal symbiont with the required elements to repopulate the bleached coral tissue (Falkowski et al. 1993; Hoegh-Guldberg 1994; Marubini and Davies 1996), and by stimulating algal photosynthesis and carbon translocation to the host (Fernandes de Barros Marangoni et al. 2020). Due to the limitation of our experimental design, we cannot determine if mortality in elevated N and N+P would have continued if *A. cervicornis* had been maintained longer in elevated nutrients without applying heat stress. However, the fact that there was no mortality in A fragments during heat (compared with 84-100% mortality in N and N+P, Fig. 2), strongly suggests that excess nutrients were the primary cause of tissue loss. We suggest that the physiological impairment of these corals due to nutrient stress was exacerbated at elevated temperatures (Wiedenmann et al. 2013; Burkepile et al. 2019), as indicated by the larger declines in *F*_*v*_*/F*_*m*_ during heat stress in *A. cervicornis* fragments pre-exposed to N and N+P compared to A (Fig. 3).

Based on our findings, *A. cervicornis* survivorship can be significantly increased, across a range of temperatures (including temperatures ∼1°C above the local summer maximum), by improving water quality. These results offer some hope, as well as additional challenges, for coral reef conservation and restoration of *A. cervicornis*. On the one hand, coastal pollution from both point and non-point sources can be more directly addressed by local policy and management practices, while reductions in carbon emissions require global action and involve considerable time before benefits will be experienced, due to committed warming from greenhouse pollutants already in the atmosphere (Donner 2009). On the other hand, if nutrient loads are not controlled, excess nitrogen (with or without additional phosphate) could impair *A. cervicornis* survivorship (Fig. 2), exacerbating the effects of heat stress, and jeopardizing the effectiveness of recovery efforts for this iconic species.

To date, monitoring of outplanted *A. cervicornis* in Florida has shown satisfactory survivorship after one year (Schopmeyer et al., 2017; O’Donnell et al., 2018), but survivorship significantly declines after two years (Ware et al. 2020) and this may be due in part to nutrient pollution. In addition to reducing survivorship, nutrient loads could reduce *A. cervicornis* growth rates (Renegar and Riegl 2005), further hindering the recovery of coral cover by this species. The data presented here suggest that, at least for *A. cervicornis* in Florida, long-term restoration success may be impaired if water quality issues are not addressed (Ware et al., 2020). However, other species, such as *O. faveolata* and *S. siderea*, may be less susceptible to these concerns (see next section).

Finally, heat stress alone (3 weeks at ∼31°C) did not cause *A. cervicornis* mortality in our study, but significant sublethal effects were detected, such as reductions in *F*_*v*_*/F*_*m*_ (Fig. 3), chlorophyll-*a* concentration (Fig. 4), and symbiont density (Fig. 5). Indeed, declines in photochemical efficiency in *A. cervicornis* were among the highest of all experimental corals maintained in the ambient (A) treatment, surpassed only by *S. siderea* fragments hosting *Cladocopium* C3 (Fig. 3). In this study, *A. cervicornis* fragments maintained in A survived to the end of the experiment and recovered their normal pigmentation and *F*_*v*_*/F*_*m*_ values 1-2 weeks after they were removed from heat stress. However, it is likely that exposure to additional heat stress (longer duration or higher temperatures) would have resulted in widespread *A. cervicornis* mortality. For example, *A. cervicornis* corals exposed to 32°C suffered 100% mortality after one month, while *O. faveolata* + *D. trenchii* survived for more than two months (Langdon et al. 2018).

### O. faveolata *and* S. siderea *susceptibility to heat stress varies with their symbiont community, but it is not affected by pre-exposure to elevated nutrients*

Bleaching susceptibility in scleractinian corals is known to vary as a function of algal symbiont community structure (Baker et al. 2004; Berkelmans and van Oppen 2006), relative abundance (Cunning and Baker 2013), and nutritional state (Wiedenmann et al. 2013). On Florida’s Coral Reef, *A. cervicornis* have been found in association with *Symbiodinium, Cladocopium*, and *Durusdinium* (Baums et al. 2010), but our experimental fragments only hosted the most common symbiont for this species in Florida, *S. fitti*. Conversely, our *O. faveolata* and *S. siderea* colonies hosted different algal symbiont types, which allowed us to compare the sensitivity of these different coral-algal combinations to elevated nutrients and heat stress.

We found that *O. faveolata + Breviolum* was susceptible to heat, with mortality beginning during the first week of heat stress (Fig. 2, ESM2-Fig. S4). Nutrient treatment in this holobiont did not affect survivorship probabilities (Fig. 2), *F*_*v*_*/F*_*m*_ (Fig. 3), or symbiont densities (Fig. 5), but did affect chlorophyll-*a* content, which increased by 49-55% in the N and N+P treatments (Fig. 4). In contrast, there was no significant mortality in response to heat stress in *O. faveolata* colonies hosting *D. trenchii* (ESM2-Figs. S1, S4). Although there were small changes in these corals under elevated nutrients and ambient temperatures (higher *F*_*v*_*/F*_*m*_ and chlorophyll-*a* content), their response to heat stress was not affected by pre-exposure to nutrients (Figs. 2-5).

Our experiment only included one *O. faveolata* colony hosting *Breviolum*, but our results agree with previous studies indicating higher thermotolerance of *O. faveolata* containing *Durusdinium* compared to *Breviolum* (e.g., Cunning et al. 2015, 2018; Parker et al. 2020). *O. faveolata* is remarkably flexible in its associations with Symbiodiniaceae and can host *Symbiodinium, Breviolum, Cladocopium, Durusdinium*, or a combination of all four of these genera (Rowan et al. 1997; Kemp et al. 2014; Kennedy et al. 2016). *Breviolum* was the most common and dominant symbiont in *O. faveolata* in the Florida Keys (Thornhill et al. 2006; Baums et al. 2010; Kemp et al. 2015), but more recent studies have shown a shift in dominance towards *D. trenchii* (Manzello et al. 2019). It is likely that severe back-to-back bleaching events in 2014 and 2015 resulted in higher mortality of corals hosting thermosensitive symbionts such as *Breviolum*, as well as shifts in the symbiont communities in surviving corals in favor of *D. trenchii* (Cunning et al. 2015). These changes in the algal community have potentially increased the thermotolerance of *O. faveolata* populations in the Caribbean, decreasing the risk of coral mortality during future heat stress events. However, there are likely to be tradeoffs, such as reduced growth rates, or less reproductive output that could still compromise the ecological function or inclusive fitness of these corals (Jones and Berkelmans 2010).

Similar to *O. faveolata, S. siderea* can host multiple algal partners, and commonly become dominated by *D. trenchii* during or after heat stress (Cunning et al. 2018). Algal symbionts in the genera *Symbiodinium, Breviolum, Cladocopium*, and *Durusdinium* have been reported in this species (Thornhill et al. 2006; Correa et al. 2009), but our colonies only hosted *Cladocopium* C1, *Cladocopium* C3, *D. trenchii* or a combination of *Cladocopium* and *D. trenchii*. In our experiment, the composition of the algal symbiont community had the strongest effect on *S. siderea* symbiont density (Fig. 5), photochemical efficiency (Fig. 3), and bleaching susceptibility (Figs. 4-5), but nutrients did not modify these patterns. Based on coral mortality alone, *S. siderea* was the most resistant species to both elevated nutrients and heat stress, with no significant mortality among any of the treatments or colonies (Fig. 2). However, colonies that hosted *Cladocopium* C3 had the strongest declines in *F*_*v*_*/F*_*m*_ of all the holobionts tested (Fig. 3) suggesting this was a relatively heat-sensitive combination compared to *Cladocopium* C1 and *D. trenchii* (Figs. 3-5).

This functional variation among *Cladocopium* types suggests that higher thermotolerance in *S. siderea* can be achieved by hosting *Cladocopium* C1 as well as by hosting *D. trenchii* (Figs. 3-5). *Cladocopium* C1 and C3 are both considered to be generalists since they are found in multiple hosts and have a global distribution (LaJeunesse 2005). While *Cladocopium* C3 has been generally considered to be thermosensitive, the relative thermotolerance of *Cladocopium* C1 in the Caribbean has not been determined. On the Great Barrier Reef, *Cladocopium* C1 has been found to exhibit local adaptation to thermal stress (Howells et al. 2011), but this has not yet been recorded in Caribbean corals. It is not yet clear whether the higher thermotolerance of C1 compared to C3 is also accompanied by physiological tradeoffs, such as reduced growth rates, that have been reported for hosts containing *D. trenchii* (Pettay et al. 2015).

A variety of species, including *S. siderea, Colpophyllia natans, O. annularis, O. franksi, Porites astreoides*, and *Pseudodiploria strigosa* have been found to host *Cladocopium* C1 in the Caribbean, including the US Virgin Islands and Yucatán peninsula of Mexico (LaJeunesse 2002, 2005; Correa et al. 2009; Finney et al. 2010; Cunning et al. 2017; Davies et al. 2018). However, this symbiont does not appear to have been very abundant in Florida in previous studies (Thornhill et al. 2006; Correa et al. 2009). Future studies should determine whether this symbiont imparts higher heat tolerance to other coral species in addition to *S. siderea*, what the tradeoffs may be, and whether this symbiont is becoming more common in the region in response to recurrent heat stress. Answering these questions will help determine if this symbiont might play a role in the future persistence of Caribbean corals under projected climate change scenarios.

## Supporting information

Electronic Supplemental Material ESM1

Electronic Supplemental Material ESM2

Electronic Supplemental Material ESM3

## Acknowledgments

We thank D. Lirman for providing *A. cervicornis* corals, and Mote Marine Laboratory for originally providing these genotypes to D. Lirman. We thank P. Rosen, R. Gilpin, C. Leto, and S. Diaz de Villegas for help with coral maintenance, and experimental data collection. Funding was provided by the National Science Foundation (NSF OCE-1358699 to AB), COLCIENCIAS (grant 529 to AMPC), and the David Rowland Endowed fellowship (to AMPC).

## Data availability

Temperature, mortality, *F*_*v*_*/F*_*m*_, chlorophyll-*a*, symbiont areal density, and qPCR data, as well as R code for analysis are available at Zenodo (see reference Palacio 2021).

