## Supplementary material for "Variation in susceptibility among three Caribbean coral species and their algal symbionts indicates the threatened staghorn coral, *Acropora cervicornis*, is particularly susceptible to elevated nutrients and heat stress": Electronic Supplemental Material ESM1

### Electronic Supplementary Material 1 (ESM1): Supplementary Methods for

#### *Coral collection*

After coral collection and fragmentation, all coral fragments were maintained in indoor tanks at ~ 26°C and 12:12 h light:dark schedule (~350 µmol PAR $m^{-2}$ $s^{-1}$) for at least two months before starting the experiment, allowing for recovery from handling and fragmentation. During the recovery time, some *A. cervicornis* experienced rapid tissue loss (RTL) over a 1-2 day period, which caused total mortality of fragments. This event was more severe in some genets, resulting in an unbalanced number of fragments from each genotype in the experiment (Table S1). No further mortality was observed during the following months of acclimation to the tank conditions.

#### *Experimental conditions*

Full water changes were performed every 2 days by supplying each aquarium with 30 L of filtered seawater from Bear Cut, Virginia Key, FL. To reach the treatment concentrations, N and N+P aquaria were supplied with 1 mL of NH4Cl [300 mM] solution, and N+P aquaria with 1 mL of NaH2PO4H2O [30 mM]. Additionally, peristaltic pumps replenished the nutrient concentrations through the addition of small nutrient doses to account for nutrient uptake. Pumps in the N aquaria delivered 0.2 mL of NH4Cl [6.25 mM] every 12 minutes to replenish 5 µmol of NH4 per day, while pumps in the N+P aquaria delivered 0.2mL of NH4Cl [6.25 mM] + NaH2PO4H2O [1.25 mM] every 12 minutes to replenish 5 µmol of NH4 and 1 µmol of PO4 per day.

Throughout the experiment (days 1-113), the corals were fed 3X per week with Zeigler® AP 110 dry larval diet by transferring them into a separate feeding tank. During the feeding time (~ 1 h), experimental aquaria were cleaned, seawater replaced, and nutrient levels replenished. Because light levels varied between different positions in the tanks (321 - 420 µmol PAR $m^{-2}$ $s^{-1}$), we rotated the position of treatments within the tanks as well as the replicate fragment assigned to each tank before transferring the corals back to their respective aquarium. This range of light intensity was maintained during the control, ramp-up and heat stress periods (321 - 420 µmol PAR $m^{-2}$ $s^{-1}$). The temperature was measured in each aquarium every 30 min using Hobo pendant data loggers (Model UA-002-08, Onset corporation).

#### *Algal symbiont community function (Fv/Fm)*

$F_v/F_m$ was measured using a MAXI Imaging Pulse Amplitude Modulated (IPAM) Fluorometer (Waltz, Effeltrich, Germany). All the corals were dark-adapted for 30 minutes before data collection. For *A. cervicornis,* all measurements were taken from the middle of the branches. For *O. faveolata* and *S. siderea*, we measured the whole core area, avoiding edges that may contain filamentous algae which can confound fluorometric data. The period between 9 to 10 AM was targeted for the IPAM data collection. However, this was not always possible and some of the measurements were done as late as 2 PM.

#### *Algal symbiont areal densities, and chlorophyll-*a *concentrations*

Tissue from each preserved fragment (Table S2) was removed with an airbrush and DNA buffer (10 mM Tris, 1 mM EDTA) and the resulting tissue slurry was homogenized using a tissue grinder. The total slurry volume was recorded before dividing it into aliquots for the different analyses. Aliquots of 0.5

- 1 mL were preserved with 50µL of Lugol's iodine solution for algal cell counts. Symbiont cells were counted using an inverted microscope and a hemocytometer. Each sample was counted twice and the mean value used for density calculations.

A second tissue slurry aliquot (2 - 5 mL) was filtered onto a Whatman glass microfiber GF/F filter (Schleicher & Schuell) to estimate chlorophyll-*a* concentration. The filters were immediately transferred to 15 mL Falcon tubes containing 3 - 4 mL of methanol and stored at -20°C for 24 h to extract the pigments. The next day, samples were vortexed and centrifuged until the supernatant was clear. Chlorophyll-*a* concentration was estimated in 2-3 supernatant replicates using a fluorometer (TD-700 Turner Designs) calibrated for the Lorenzen's modified monochromatic method using chlorophyll-*a* standards. To avoid chlorophyll degradation, the samples were kept on ice, covered with aluminum foil, and the readings were performed in a dark room.

Algal symbiont cell counts and chlorophyll-*a* values were corrected for any dilution introduced during the processing, and by the total volume of blastate obtained from each coral fragment. These values were then normalized to coral surface area (symbiont cells $cm^{-2}$, and µg chlorophyll-*a* $cm^{-2}$). For *A. cervicornis* we approximated the area of the fragment covered by tissue to the lateral surface of a cylinder (area = πdh), using the length of the fragment (h) and the mean value of the apical and basal diameters (d). For *O. faveolata* and *S. siderea,* the surface area of the cores was calculated from pictures that included a size scale using CPCe Software (Kohler and Gill 2006).

#### *Symbiodyniaceae community structure (qPCR assays)*

Each TaqMan qPCR reaction (10 µL) included 5.00 µL of TaqMan Master (Thermo Fisher Scientific, Waltham, MA), 0.5 µL of F and R primers at variable concentrations (Table S3), 0.5 µL of probe at variable concentrations (Table S3), 1 µL of DNA template, and water. SYBR-Green qPCR reactions included 6.25 µL of SYBR-Green (Thermo Fisher Scientific, Waltham, MA), 0.1125 µL of F primer [100 µM], 0.1125 µL of R primer [100 µM], 1 µL of DNA template, and 5.025 µL of water.

We first surveyed the Symbiodiniaceae genera present in each coral species by running qPCR reactions for a subset of samples from all colonies (n = 85 samples for *A. cervicornis*, n = 22 for *O. faveolata*, and n = 42 for *S. siderea*) in which we targeted the four main algal genera present in Caribbean scleractinian corals (*Symbiodinium, Breviolum, Cladocopium*, and *Durusdinium*). If qPCR failed to amplify a symbiont genus in every sample from one coral species, we assumed that the genus was not associated with the host and it was not included in further qPCR analysis for that coral species. Once the symbiont targets were defined for each coral species, all the DNA samples were amplified in duplicate reactions per target (host and algal symbionts), and the cycle threshold ($C_T$) values in which the fluorescent signal crossed the $\Delta R_n$ threshold (Table S3) were obtained from the qPCR machine. Amplifications were quality controlled by removing (1) data from plates with amplification in no-template (negative) controls, (2) amplifications in which only one of the two technical replicates crossed the fluorescence threshold, and (3) samples in which the $C_T$ standard deviation between technical replicates was higher than 1.5.

S/H cell ratios were estimated using the StepOne package for R (Cunning 2018). This package averages $C_T$s among technical replicates and corrects them with supplied information about target ploidy [coral host = 2, Symbiodiniaceae = 1 (Santos and Coffroth 2003)]; DNA extraction efficiency [coral host = 0.982, Symbiodiniaceae = 0.813 (Cunning and Baker 2013)], and gene copy number [*A. cervicornis* (CAM) = 1 (Schwartz et al. 2012), *O. faveolata* (SC1) = 1 (Severance et al. 2004), *S. siderea* (Pax-C) = 1 (van Oppen et al. 2000), *Symbiodinium* (Actin) = 9 (See methods below), *Breviolum* (Actin) = 1 (Cunning et al. 2015), *Cladocopium* (Actin) = 23, and *D. trenchii* (Actin) = 2 (Cunning 2013)]. All the primers used have amplification efficiency close to 100% and therefore can be used to compare the different targets (Cunning and Baker 2013; Cunning et al. 2015, methods below). Fluorometry corrections were done among the different TaqMan assays (Symbiodiniaceae genus), but not among host and symbiont targets run with SYBR-Green and TaqMan Master Mix, respectively. Therefore S/H cell ratios for *O. faveolata* and *S. siderea* are relative ratios and can be used to compare their relative changes in symbiont abundance among nutrient or heat treatments in each species, but not to compare symbiont abundance among different coral species. The total symbiont to host (S/H) cell ratio was calculated for the corals that hosted multiple Symbiodiniaceae genera as the addition of all genus cell ratios (S/H = *Symbiodinium*/Host + *Breviolum*/Host + *Cladocopium*/Host + *Durusdinium*/Host). For *O. faveolata* and *S. siderea*, we calculated the proportion of the symbiont community composed by thermotolerant *D. trenchii* as (*D.trenchii*/Host)/(S/H).

##### *Single copy qPCR assay development for Caribbean* **Acropora**

Available DNA sequences from the single-copy genes Pax-C and Calmodulin (CaM) in the Caribbean *Acropora* spp. were obtained from (Schwartz et al. 2012; Table S4) and were manually aligned using Unipro *EUGene*®. We used the resulting consensus alignment for each locus to search for candidate primers with PrimerExpress®. The best candidates (Pax-C forward: 5'- TTGTCACTTTTA CGCGCCTAGA-3', reverse: 5'-GCAGTGCACGCTCTTCTTCTC - 3'; and CaM forward: 5'- GCC CTAATTTCTGATCGATTCAA-3', reverse: 5'-GCAGACAGAAGGGCCACT-3') were synthesized by Integrated DNA Technologies Inc, and their amplification efficiency was compared using the standard curve method. First, we created serial dilutions of *A. cervicornis* genomic DNA extracted from five coral colonies, and amplified each dilution by triplicate with both sets of primers. Reaction volumes were 12.5 µL, with 6.25 µL SYBR Green, 0.1125 µL F primer [100 µM], 0.1125 µL R primer [100 µM], 5.025 µL water and 1 µL DNA template. Amplification efficiency was calculated from the slope of the standard curve ($C_T$ vs log10[dilution factor]) using the equation Efficiency = [$10^{-1/slope}$) - 1]*100. Linear regressions estimated for both set of primers showed high amplification efficiency (Pax-C = 106.9% ± 4.0 SD, CaM = 105.2% ± 4.7 SD), and therefore can be compared with other validated Symbiodiniaceae assays to calculate S/H cell ratios in *A. cervicornis* (Table S5). CaM assay was chosen over Pax-C because its amplification efficiency was slightly closer to 100%.

##### *Symbiodiniaceae actin gene copy number*

*Symbiodinium fitti* actin copy number estimation followed the method described in Cunning and Baker (2013). Briefly, coral tissue slurries with a known *Symbiodinium* density (cells $\mu L^{-1}$; see methods for Symbiodiniaceae areal density estimation) were obtained from three *A. cervicornis* fragments. We extracted the DNA from five replicate aliquots per sample, each one estimated to contain 100,000 *Symbiodinium* cells. These DNA samples were then quantified using qPCR standard curves of copy number standards, assuming a 95% extraction efficiency for the slurry samples. The calculated actin locus copy number for *Symbiodinium* was 8.7± 0.3 and therefore a copy number of nine was used to correct the A/H cell ratios. Actin copy number of 1 was used for *Breviolum* associated with *O. faveolata* following Cunning et al. (2015), 23 for *Cladocopium* associated with *S. siderea*, and three for *Durusdinium* associated with *O. faveolata* and *S. siderea* (Cunning 2013).

#### *Molecular identification of* **Cladocopium** *taxa*

A subset of *S. siderea* samples that hosted *Cladocopium* were amplified in PCR reactions using the primers 'ITS 2 clamp' and 'ITSintfor2' following LaJeunesse and Trench (2000). PCR products were run in denaturing gradient gel electrophoresis (DGGE) (45–80%) for 15 hours at 96 volts and 60°C (CBS Scientific, San Diego, CA) following LaJeunesse (2002) and the dominant DGGE bands in each ITS2 fingerprint were excised and re-amplified using the primers 'ITS 2' (without clamp) and 'ITSintfor2' (LaJeunesse and Trench 2000). Amplified products were purified and sequenced with their respective reverse and forward primers. Reverse and forward chromatograms were manually checked and aligned using Geneious Prime 2020.0.3.

**Table S1:** Number of fragments exposed to each nutrient treatment per coral species and genotype/colony. For coral species *A. cer* = *Acropora cervicornis*, *O. fav* = *Orbicella faveolata*, and *S. sid* = *Siderastrea siderea*. For the algal symbionts A = *Symbiodinium*, B = *Breviolum*, C = *Cladocopium* and D = *Durusdinium.*

| **Coral species** | **Coral colony + (Symbiodiniaceae genera hosted)** | **Nutrient treatment / Replicate tank (R)** | | | | | | **Total** |
|---|---|---|---|---|---|---|---|---|
| | | **A** (ambient nutrients) | | **N** [ambient nutrients + 10µM $NH_4$] | | **N+P** [ambient nutrients + 10µM $NH_4$ + 1µM $PO_4$] | | |
| | | **R1** | **R2** | **R1** | **R2** | **R1** | **R2** | |
| *A. cer* | G_48 (A) | 5 | 5 | 5 | 4 | 5 | 4 | **28** |
| | G_62 (A) | 5 | 4 | 5 | 5 | 5 | 5 | **29** |
| | G_31 (A) | 3 | 2 | 3 | 3 | 3 | 2 | **16** |
| | G_07 (A) | 5 | 4 | 5 | 4 | 4 | 4 | **26** |
| | G_50 (A) | 2 | 2 | 2 | 2 | 3 | 2 | **13** |
| | G_08 (A) | 1 | 1 | 1 | 2 | 1 | 2 | **8** |
| | **Total *A. cer*** | **21** | **18** | **21** | **20** | **21** | **19** | **120** |
| *O. fav* | Of_34 (D) | 3 | 3 | 3 | 3 | 3 | 3 | **18** |
| | Of_20 (D) | 3 | 3 | 3 | 3 | 3 | 3 | **18** |
| | Of_6 (D) | 3 | 3 | 3 | 3 | 3 | 3 | **18** |
| | Of_31 (B>>D) | 3 | 3 | 3 | 3 | 3 | 3 | **18** |
| | **Total *O. fav*** | **12** | **12** | **12** | **12** | **12** | **12** | **72** |
| *S. sid* | Ss_22 (D) | 4 | 4 | 4 | 4 | 4 | 4 | **24** |
| | Ss_23 (D) | 4 | 4 | 4 | 4 | 4 | 4 | **24** |
| | Ss_27 (D) | 4 | 4 | 4 | 4 | 4 | 4 | **24** |
| | Ss_28 C1≈D | 3 | 4 | 3 | 4 | 3 | 3 | **20** |
| | Ss_20 (C1) | 4 | 4 | 4 | 4 | 4 | 4 | **24** |
| | Ss_24 (C3>D) | 4 | 4 | 4 | 4 | 4 | 4 | **24** |
| | Ss_30 (C3) | 4 | 4 | 4 | 4 | 4 | 4 | **24** |
| | **Total *S. sid*** | **27** | **28** | **27** | **28** | **27** | **27** | **164** |

**Table S2:** Number of whole fragment samples collected to estimate symbiont areal densities and Chlorophyll-*a* concentration per nutrient treatment, temperature phase, and coral species. A: Ambient nutrients, N: elevated $NH_4$ by 10µM, and N+P: elevated $NH_4$ by 10µM + and $PO_4$ by 1µM.

| Coral Species | Temperature phase | Nutrient treatment | | | |
|---|---|---|---|---|---|
| | | A | N | N+P | Total |
| *A. cervicornis* | Control | 9 | 9 | 10 | **28** |
| | Heat | 10 | 0 | 1 | **11** |
| *O. faveolata* | Control | 8 | 8 | 8 | **24** |
| | Heat | 6 | 6 | 5 | **17** |
| *S. siderea* | Control | 14 | 14 | 13 | **41** |
| | Heat | 14 | 14 | 14 | **42** |
| **Total** | | **61** | **52** | **51** | **163** |

**Table S3:** Reaction conditions for qPCR assays used to estimate the symbiont to host (S/H) cell ratio in each coral host. *A. cervicornis* (*A.cer*) corals were only found in association with *Symbiodinium* (clade A) (yellow assays). *O. faveolata (O.fav)* was found in association with *Breviolum* (clade B), *Cladocopium* (clade C), and *Durusdinium* (clade D, red assays). *S. siderea (S.sid)* hosted *Cladocopium* and *Durusdinium* (purple assays).

| **Target** | **A** | ***A. cer*** | ***O. fav*** | **B** | **C** | **D** | ***S. sid*** |
|---|---|---|---|---|---|---|---|
| **Type of assay** | SYBR Green<br>Singleplex | SYBR Green<br>Singleplex | SYBR Green<br>Singleplex | TaqMan<br>Singleplex | TaqMan<br>Multiplex | | SYBR Green<br>Singleplex |
| **Primers target** | Actin | CaM | SC_Ofav | Actin | Actin | Actin | Pax-C |
| **Forward Primer** | 900nM | 900nM | 900nM | 200 nM | 50nM | 50nM | 900nM |
| **Reverse Primer** | 900nM | 900nM | 900nM | 300 nM | 75nM | 75nM | 900nM |
| **Probe** | NA | NA | NA | 100nM (FAM) | 100nM (VIC) | 100nM (FAM) | NA |
| **MasterMix** | 6.25 µl | 6.25 µl | 6.25 µl | 5 µl | 5 µl | | 6.25 µl |
| **DNA template** | 1 µl | 1 µl | 1 µl | 1 µl | 1 µl | 1 µl | 1 µl |
| **Total volume** | 12.5 µL | 12.5 µL | 12.5 µL | 10 µL | 10 µL | | 12.5 µL |
| **Machine used** | Quant Studio3 | Quant Studio3 | StepOne Plus | StepOne Plus | StepOne Plus | | StepOne Plus |
| **$\Delta R_n$ or CT threshold** | 0.2 | 0.2 | 0.2 | 0.02 | 0.02 | | 0.2 |
| **Assay reference** | (Winter 2017) | New (See below) | Cunning et al. (2015) | | Cunning and Baker (2013) | | Cunning, R. (2013) |

**Table S4:** GenBank accession numbers for the DNA sequences from (Schwartz et al. 2012)) used to develop single copy qPCR assays for the Caribbean *Acropora* spp.

| Loci | *A. cervicornis* | *A. palmata* |
|---|---|---|
| Pax-C | EU918781.1, EU918780.1<br>JN871694.1, JN871693.1<br>JN871692.1, JN871691.1<br>AF344356.1, AF344355.1 | EU918783.1, EU918782.1,<br>AF344412.1, AF344411.1 |
| Calmodulin (CaM) | EU534132.1, EU534131.1<br>EU534130.1, EU534129.1<br>EU534128.1, EU534127.1 | EU534140.1, EU534139.1<br>EU534138.1, EU534137.1<br>EU534136.1, EU534135.1<br>EU534134.1, EU534133.1 |

**Table S5:** Amplification efficiency for *A. cervicornis* primers used in new qPCR assays. Primers were tested in serial dilutions from five different coral genotypes donated from the MOTE coral nursery in the Florida Keys.

| Locus | Coral sample | Intercept | Slope | r squared | % Efficiency |
|---|---|---|---|---|---|
| Pax-C | Acer 105 | 19.623 | -3.306 | 0.992 | 100.683 |
| | Acer 116 | 20.715 | -3.157 | 0.976 | 107.367 |
| | Acer 149 | 19.670 | -3.187 | 0.991 | 105.976 |
| | Acer 162 | 19.438 | -3.163 | 0.970 | 107.079 |
| | Acer 176 | 20.707 | -3.041 | 0.983 | 113.212 |
| | **Mean** | **20.031** | **-3.171** | **0.983** | **106.864** |
| CaM | Acer 105 | 20.457 | -3.092 | 0.968 | 110.571 |
| | Acer 116 | 20.862 | -3.252 | 0.979 | 103.021 |
| | Acer 149 | 19.785 | -3.396 | 0.995 | 97.004 |
| | Acer 162 | 19.803 | -3.153 | 0.989 | 107.557 |
| | Acer 176 | 20.631 | -3.150 | 0.986 | 107.695 |
| | **Mean** | **20.307** | **-3.209** | **0.983** | **105.114** |

#### ***Cited Literature***

Cunning R (2013) The role of algal symbiont community dynamics in reef coral responses to global climate change. Ph.D. thesis, University of Miami, FL, USA

Cunning R (2018) SteponeR: R package for importing qPCR data from StepOne™ Software

Cunning R, Baker AC (2013) Excess algal symbionts increase the susceptibility of reef corals to bleaching. Nat Clim Chang 3:259

Cunning R, Silverstein RN, Baker AC (2015) Investigating the causes and consequences of symbiont shuffling in a multi-partner reef coral symbiosis under environmental change. Proc R Soc Lond B Biol Sci 282:20141725

Kohler KE, Gill SM (2006) Coral Point Count with Excel extensions (CPCe): A Visual Basic program for the determination of coral and substrate coverage using random point count methodology. Comput Geosci 32:1259–1269

LaJeunesse TC (2002) Diversity and community structure of symbiotic dinoflagellates from Caribbean coral reefs. Mar Biol 141:387–400

LaJeunesse TC, Trench RK (2000) Biogeography of two species of *Symbiodinium* (Freudenthal) inhabiting the intertidal sea anemone *Anthopleura elegantissima* (Brandt). Biol Bull 199:126–134

van Oppen MJH, Willis BL, Van Vugt HWJA, Miller DJ (2000) Examination of species boundaries in the *Acropora cervicornis* group (Scleractinia, Cnidaria) using nuclear DNA sequence analyses. Mol Ecol 9:1363–1373

Santos SR, Coffroth MA (2003) Molecular genetic evidence that dinoflagellates belonging to the genus *Symbiodinium* freudenthal are haploid. Biol Bull 204:10–20

Schwartz SA, Budd AF, Carlon DB (2012) Molecules and fossils reveal punctuated diversification in Caribbean “faviid” corals. BMC Evol Biol 12:123

Severance EG, Szmant AM, Karl SA (2004) Single-copy gene markers isolated from the Caribbean coral, *Montastraea annularis*. Mol Ecol Notes 4:167–169

Winter RN (2017) Environmental Controls on the Reassembly of *Symbiodinium* Communities in Reef Corals Following Perturbation: Implications for Reef Futures Under Climate Change. Ph.D. thesis, University of Miami, FL, USA
