## Supplementary material for "Variation in susceptibility among three Caribbean coral species and their algal symbionts indicates the threatened staghorn coral, *Acropora cervicornis*, is particularly susceptible to elevated nutrients and heat stress": Electronic Supplemental Material ESM2

### Electronic Supplementary Material 2 (ESM2):
### Supplementary Figures for

#### Variation in susceptibility among three Caribbean coral species and their algal symbionts indicates the threatened *Acropora cervicornis* is particularly susceptible to elevated nutrients and heat stress

Ana M. Palacio-Castro, Caroline E. Dennison, Stephanie M. Rosales, Andrew C. Baker

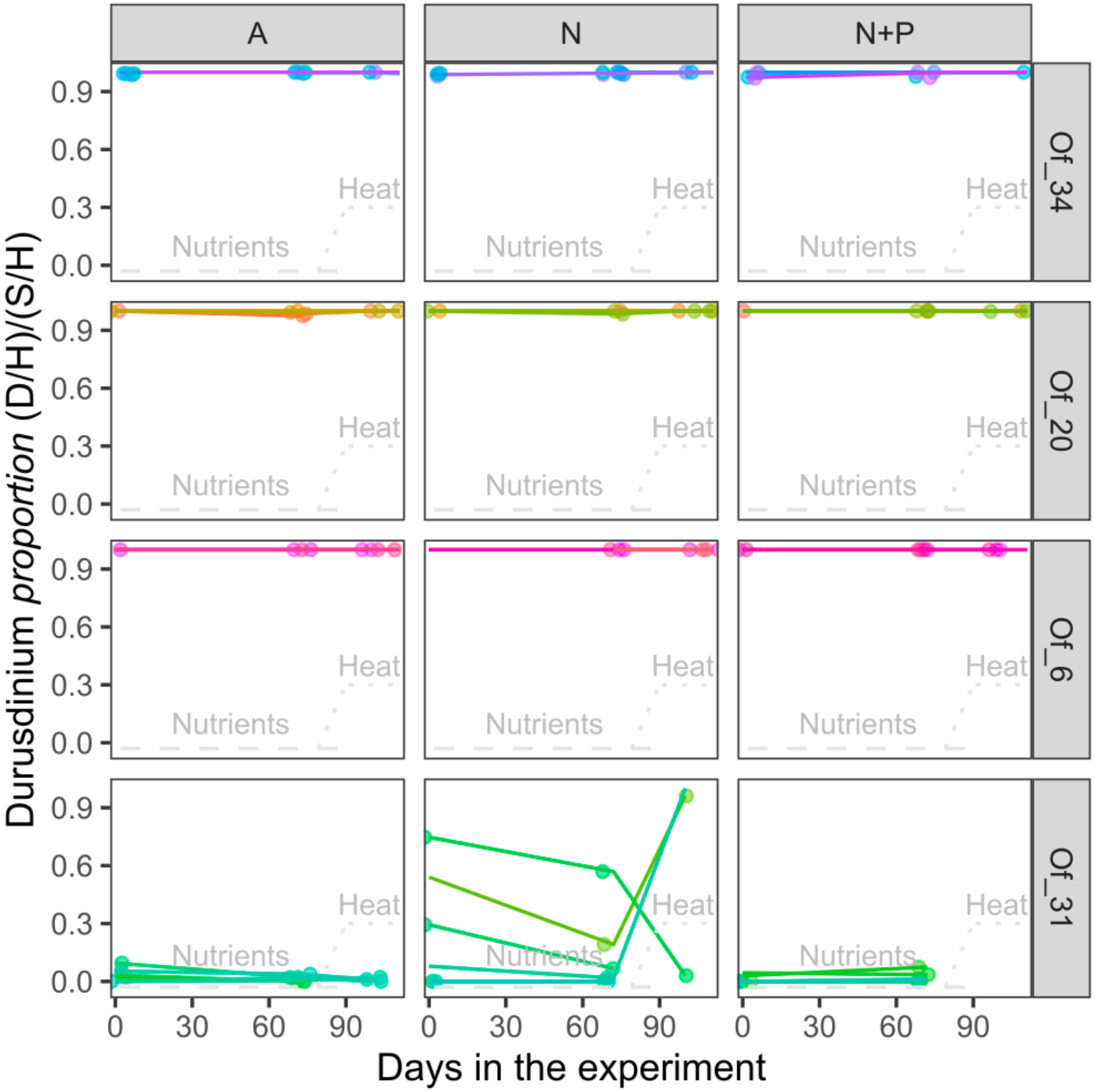


**Figure S1: Proportion of *Durusdinium* in the total symbiont community of *O. faveolata* cores through the experiment.** Each panel groups the cores from an individual colony under each nutrient treatment (A, N, and N+P) during control, ramp-up, and heat stress phases. Colonies Of_34, Of_20 and Of_6 were dominated by *D. trenchii.* Colony Of_31 was mainly dominated by *Breviolum*, but some cores hosted variable amounts of *D. trenchii* and *Breviolum.*

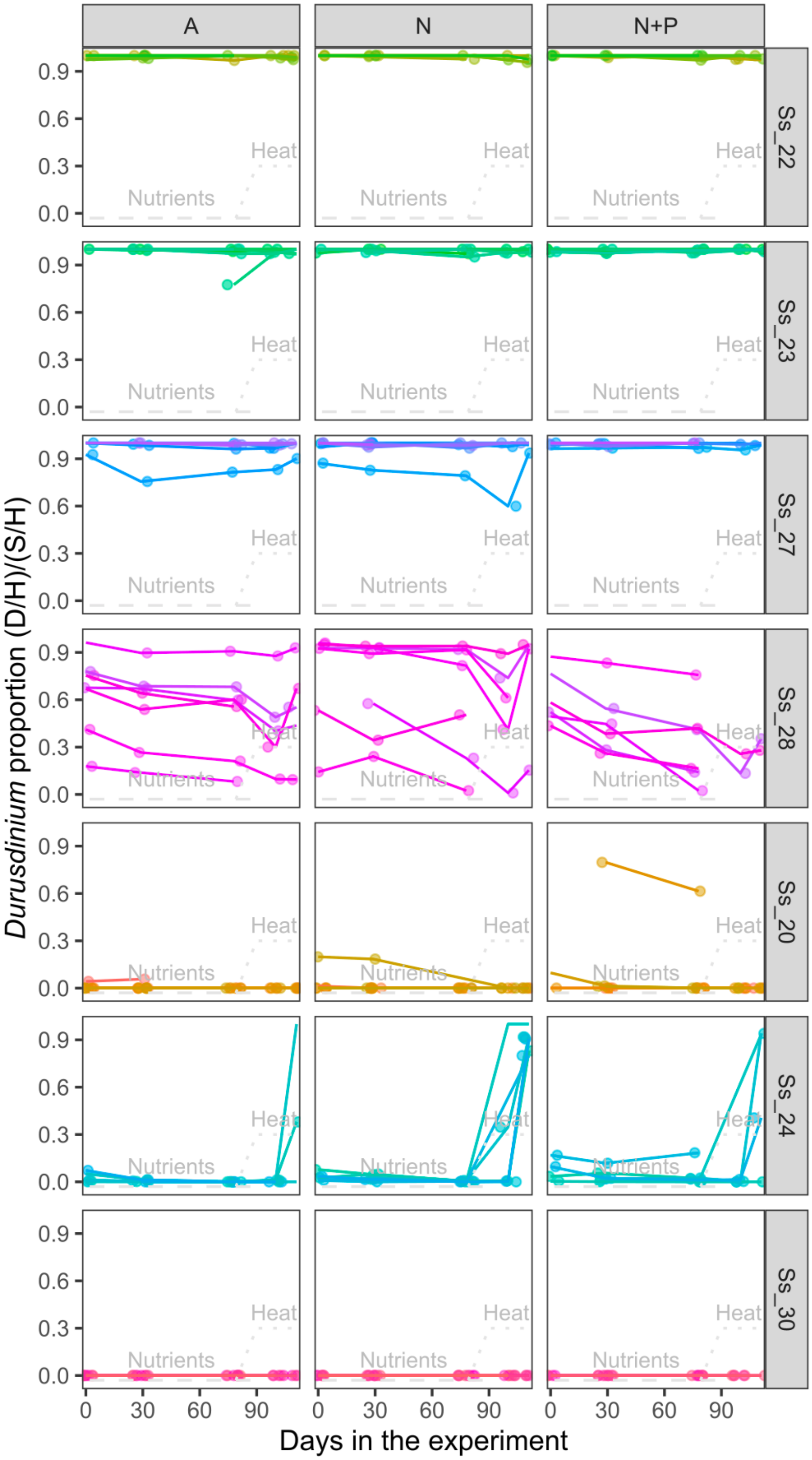


**Figure S2: Proportion of *Durusdinium* in the total symbiont community of *S. siderea* cores through the experiment**. Each panel groups cores from an individual colony under each nutrient treatment (A, N, and N+P) during control, ramp-up, and heat stress phases. Colonies Ss_22, Ss_23 and Ss_27 were dominated by *D. trenchii.* Colony Ss_28 hosted variable amounts of *D. trenchii* and *Cladocopium* C1. Colony Ss_20 was dominated by *Cladocopium* C1. Colonies Ss_24 and Ss_30 were initially dominated by *Cladocopium* C3, but Ss_24 had background *D. trenchii* and became dominated by this last symbiont during heat stress.

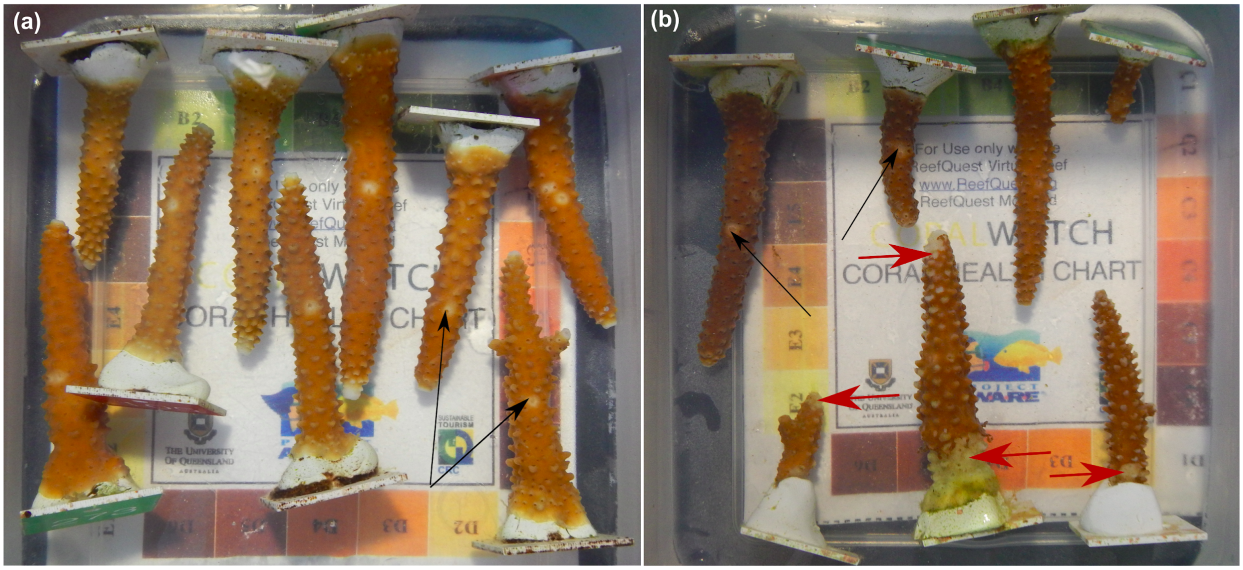


**Figure S3: Example of fragments presenting rapid tissue loss (RTL at day 84).** Black arrows show examples of normal tissue scars due to biopsy samples. Red arrows show areas of the fragments experiencing RTL. **(a)** Healthy fragments maintained in ambient nutrients **(b)** Fragments maintained in elevated N+P, some of them showing RTL.

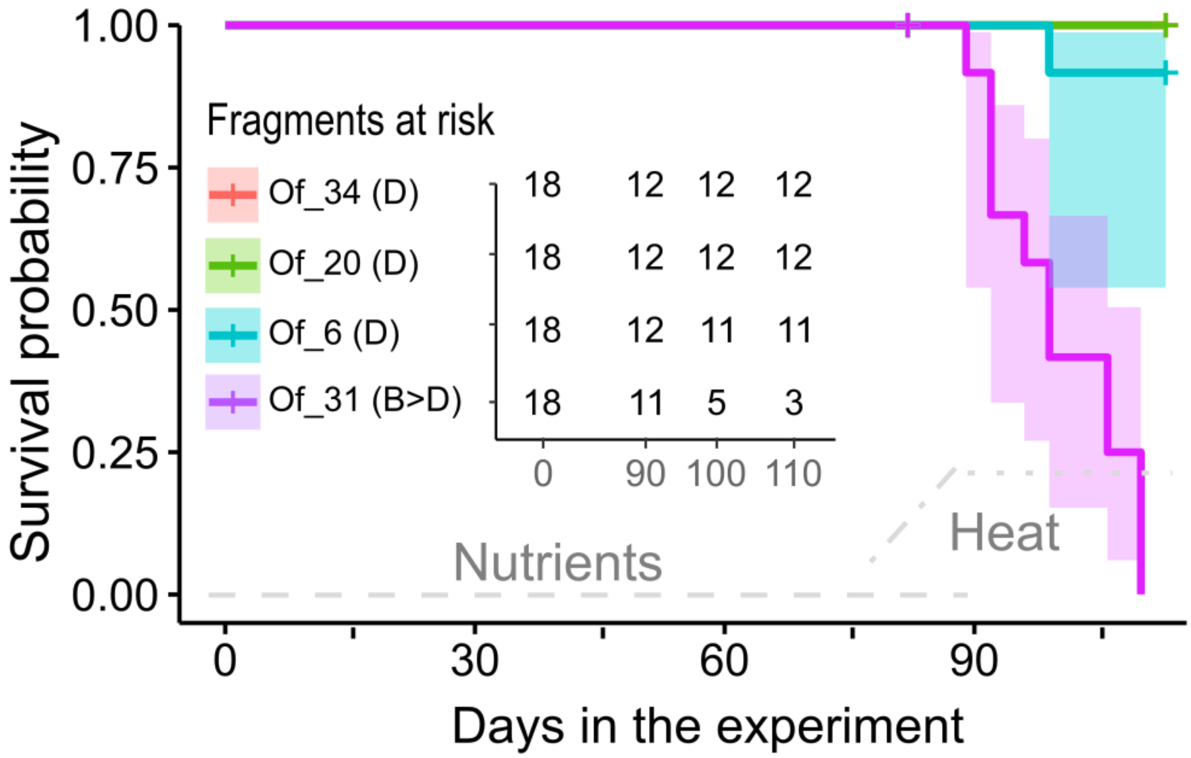


**Figure S4: Differential effect of heat stress on *O. faveolata* survivorship**. Fragments at risk tables show the number of fragments from each colony that were in the experiment at any specific day. The "+" in the graphs represent "censored" events in which corals were removed from the tanks to obtain whole-fragment samples and therefore are excluded from further survivorship analysis. *O. faveolata* survivorship was not affected by the addition of nutrients ($p = 0.85$), but heat stress significantly reduced survivorship probabilities in one colony that hosted *Breviolum* (B) instead of *D. trenchii* (D) ($p < 0.0001$).
