## Supplementary material for "Variation in susceptibility among three Caribbean coral species and their algal symbionts indicates the threatened staghorn coral, *Acropora cervicornis*, is particularly susceptible to elevated nutrients and heat stress": Electronic Supplemental Material ESM3

### Electronic Supplementary Material 3 (ESM3):
### Model outputs for

**Variation in susceptibility among three Caribbean coral species and their algal symbionts indicates the threatened *Acropora cervicornis* is particularly susceptible to elevated nutrients and heat stress**

Ana M. Palacio-Castro, Caroline E. Dennison, Stephanie M. Rosales, Andrew C. Baker

This PDF contains:


- Mixed model summaries
    - Table S6 (*Fv/Fm* - all coral species)
    - Table S7 (Chlorophyll-*a* - all coral species)
    - Table S8 (Symbiodiniaceae areal densities - all coral species)

- Tukey HSD tests and variable relative changes
    - Table S9 (*A. cervicornis Fv/Fm* )
    - Table S10 (*O. faveolata Fv/Fm* )
    - Table S11 (*S. siderea Fv/Fm* )
    - Table S12 (Chlorophyll-*a* - all coral species)
    - Table S13 (Symbiodiniaceae areal densities - all coral species)
    - Table S14 ( *S. siderea* Chlorophyll-*a* Symbiodiniaceae areal densities by Symbiodiniaceae type)

**Table S6:** Generalized linear mixed models used to test for differences in the photochemical efficiency ($F_v$ /$F_m$) of corals exposed to nutrient treatments (A, N and N+P) at control temperature (days 1-78) and heat stress (days 90-113). Mixed-effects models were run with the lme4 1.1-17 package (Bates et al. 2015) for R 3.4.3 (R Core Team 2018). Each model included *nutrient* treatment, dominant *symbiont* type (except for *A. cervicornis* that hosted only one type), and *days* in the experiment as interacting fixed factors, as well as coral *genotype*, *fragment*, and *replicate* tank as random effects.

| ***A. cervicornis* $F_v/F_m$** | | | | |
|---|---|---|---|---|
| **Fixed effects** | **numDF** | **denDF** | **F-value** | **p-value** |
| Nutrient | 2 | 120.5 | 19.6 | <0.001 |
| Days | 16 | 1460.5 | 666.7 | <0.001 |
| Nutrient:Day | 31 | 1460.6 | 69.3 | <0.001 |
| **Random effects** | **npar** | **logLik** | **AIC** | **Pr(>Chisq)** |
| Genotype | 53 | 3541.1 | -6899.2 | <0.001 |
| Fragment | 53 | 3465.9 | -6825.8 | <0.001 |
| Replicate (Tank) | 53 | 3502.6 | -6974.8 | 0.080 |
| ***O. faveolata* $F_v/F_m$** | | | | |
| **Fixed effects** | **numDF** | **denDF** | **F-value** | **p-value** |
| Nutrient | 2 | 62.7 | 2.8 | 0.07 |
| Day | 8 | 50.5 | 50.5 | <0.001 |
| Symbiont | 1 | 35.6 | 10.8 | <0.001 |
| Nutrient : Day | 16 | 528 | 4.7 | <0.001 |
| Nutrient : Symbiont | 2 | 62.7 | 1.3 | 0.27 |
| Day: Symbiont | 8 | 528 | 7.6 | <0.001 |
| Nutrient: Day: Symbiont | 16 | 528 | 2.8 | <0.001 |
| **Random effects** | **npar** | **logLik** | **AIC** | **Pr(>Chisq)** |
| Genotype | 57 | 1302.2 | -2490.5 | <0.001 |
| Fragment | 57 | 1273.0 | -2432.0 | <0.001 |
| Replicate (Tank) | 57 | 1311.5 | -2509.1 | 1 |
| ***S. siderea* $F_v/F_m$** | | | | |
| **Fixed effects** | **numDF** | **denDF** | **F-value** | **p-value** |
| Nutrient | 2 | 146.6 | 16.2 | <0.001 |
| Day | 8 | 1220.3 | 261.5 | <0.001 |
| Symbiont | 2 | 8.5 | 1.4 | 0.3 |
| Nutrient : Day | 16 | 1220.3 | 12.9 | <0.001 |
| Nutrient : Symbiont | 4 | 146.6 | 9.9 | <0.001 |
| Day: Symbiont | 16 | 1220.3 | 5.2 | <0.001 |
| Nutrient: Day: Symbiont | 32 | 1220.3 | 1.2 | 0.2 |
| **Random effects** | **npar** | **logLik** | **AIC** | **Pr(>Chisq)** |
| Genotype | 84 | 2995.3 | -5822.7 | <0.001 |
| Fragment | 84 | 2975.1 | 5782.1 | <0.001 |
| Replicate (Tank) | 84 | 3033.3 | -5898.6 | 1 |

**Table S7:** Generalized linear mixed models used to test for differences in the Chlorophyll-*a* concentration in corals exposed to nutrient treatments (A, N and N+P) at control temperature (day 78) and heat stress (day 113). Mixed-effects models were run with the lme4 1.1-17 package (Bates et al. 2015) for R 3.4.3 (R Core Team 2018). Each model included *nutrient* treatment, dominant *symbiont* type (except for *A. cervicornis* that hosted only one type), and *day* in the experiment as interacting fixed factors, as well as coral *genotype*, and *replicate* tank as random effects.

| ***A. cervicornis* Chlorophyll-*a*** | | | | |
|---|---|---|---|---|
| **Fixed effects** | **numDF** | **denDF** | **F-value** | **p-value** |
| Nutrient | 2 | 26 | 1.72 | 0.2 |
| Day | 1 | 26 | 29.5 | <0.001 |
| Nutrient:Day | 1 | 26 | 3.5 | <0.05 |
| **Random effects** | **npar** | **logLik** | **AIC** | **Pr(>Chisq)** |
| Genotype | 7 | -42.2 | 98.4 | 1 |
| Replicate (Tank) | 7 | -42.2 | 98.4 | 1 |
| ***O. faveolata* Chlorophyll-*a*** | | | | |
| **Fixed effects** | **numDF** | **denDF** | **F-value** | **p-value** |
| Nutrient | 2 | 30.0 | 2.9 | 0.06 |
| Day | 1 | 30.0 | 71.5 | <0.001 |
| Symbiont | 1 | 2.3 | 1.0 | 0.4 |
| Nutrient : Day | 2 | 30.0 | 5.4 | 0.01 |
| Nutrient : Symbiont | 2 | 30.0 | 0.3 | 0.7 |
| Day: Symbiont | 0 | NA | NA | NA |
| Nutrient: Day: Symbiont | 0 | NA | NA | NA |
| **Random effects** | **npar** | **logLik** | **AIC** | **Pr(>Chisq)** |
| Genotype | 10 | 0.3 | 19.4 | <0.001 |
| Replicate (Tank) | 11 | 7.9 | 6.1 | 0.9 |
| ***S. siderea* Chlorophyll-*a*** | | | | |
| **Fixed effects** | **numDF** | **denDF** | **F-value** | **p-value** |
| Nutrient | 2 | 57.6 | 6.2 | <0.001 |
| Day | 1 | 57.4 | 216.5 | <0.001 |
| Symbiont | 2 | 31.8 | 2.6 | 0.09 |
| Nutrient : Day | 2 | 57.9 | 2.2 | 0.1 |
| Nutrient : Symbiont | 4 | 57.3 | 0.4 | 0.8 |
| Day: Symbiont | 2 | 57.2 | 9.7 | <0.001 |
| Nutrient: Day: Symbiont | 4 | 57.6 | 0.6 | 0.7 |
| **Random effects** | **npar** | **logLik** | **AIC** | **Pr(>Chisq)** |
| none | 21 | -110.2 | 262.4 | |
| Genotype | 20 | -119.9 | 279.8 | <0.001 |
| Replicate (Tank) | 20 | -110.2 | 260.4 | 1 |

**Table S8:** Generalized linear mixed models used to test for differences in the Symbiodiniaceae areal density (cells cm$^{-2}$) in corals exposed to nutrient treatments (A, N and N+P) at control temperature (day 78) and heat stress (day 113). Mixed-effects models were run with the lme4 1.1-17 package (Bates et al. 2015) for R 3.4.3 (R Core Team 2018). Each model included *nutrient* treatment, dominant *symbiont* type (except for *A. cervicornis* that hosted only one type), and *day* (temperature) in the experiment as interacting fixed factors, as well as coral *genotype*, and *replicate* tank as random effects.

| ***A. cervicornis Symbiodiniaceae* density** | | | | |
|---|---|---|---|---|
| **Fixed effects** | **numDF** | **denDF** | **F-value** | **p-value** |
| Nutrient | 2 | 23.8 | 0.14 | 0.87 |
| Day | 1 | 24.3 | 55.5 | <0.001 |
| Nutrient : Day | 1 | 24.2 | 3.4 | 0.08 |
| **Random effects** | **npar** | **logLik** | **AIC** | **Pr(>Chisq)** |
| Genotype | 7 | -43.2 | 100.3 | 0.07 |
| Replicate (Tank) | 7 | -41.8 | 97.6 | 0.5 |
| ***O. faveolata Symbiodiniaceae* density** | | | | |
| **Fixed effects** | **numDF** | **denDF** | **F-value** | **p-value** |
| Nutrient | 2 | 30.0 | 1.0 | 0.4 |
| Day | 1 | 30.0 | 125.4 | <0.001 |
| Symbiont | 1 | 2.2 | 0.2 | 0.7 |
| Nutrient : Day | 2 | 30.0 | 4.3 | 0.02 |
| Nutrient : Symbiont | 2 | 30.0 | 1.7 | 0.20 |
| **Random effects** | **npar** | **logLik** | **AIC** | **Pr(>Chisq)** |
| Genotype | 10 | 0.3 | 19.4 | <0.001 |
| Replicate (Tank) | 11 | 7.9 | 6.1 | 0.9 |
| ***S. siderea Symbiodiniaceae* density** | | | | |
| **Fixed effects** | **numDF** | **denDF** | **F-value** | **p-value** |
| Nutrient | 2 | 58.6 | 1.8 | 0.2 |
| Day | 1 | 57.9 | 231.4 | <0.001 |
| Symbiont | 1 | 8.2 | 0.3 | 0.7 |
| Nutrient : Day | 1 | 59.6 | 0.02 | 0.9 |
| Nutrient : Symbiont | 4 | 58.7 | 0.8 | 0.5 |
| Day: Symbiont | 2 | 58.1 | 27.3 | <0.001 |
| Nutrient: Day: Symbiont | 4 | 59.2 | 0.2 | 0.9 |
| **Random effects** | **npar** | **logLik** | **AIC** | **Pr(>Chisq)** |
| none | 21 | 17.1 | 7.7 | |
| Genotype | 20 | 15.4 | 9.2 | 0.06 |
| Replicate (Tank) | 20 | 17.1 | 5.7 | 1 |

**Table S9:** Estimated photochemical efficiency ($F_v$ /$F_m$) for *A. cervicornis* corals exposed to different nutrient treatments and subsequent heat stress. Pairwise comparisons between groups were obtained using Tukey's HSD test (α = 0.05). Stars in *N and *N+P denote the group of corals that were assigned to these treatments, but that were not exposed to elevated nutrients at the time of the measurement. Percentages of change in bold represent comparison among values that were significantly different based on the Tukey's HSD test.

| ***Days* in the experiment (Phase)** | ***Nutrient* Treatment** | **$F_v/F_m$ Em mean** | **df** | **Lower CL** | **Upper CL** | **Tukey Group** | **% change respect ambient (same day)** | **% change respect baseline (Day 1)** | **% change respect control (Day 76)** |
|---|---|---|---|---|---|---|---|---|---|
| 1 (Baseline) | A | 0.61 | 10.28 | 0.60 | 0.63 | G-L | NA | NA | NA |
| | *N | 0.60 | 9.91 | 0.58 | 0.62 | E-H | -1.8% | NA | NA |
| | *N+P | 0.60 | 10.06 | 0.59 | 0.62 | G-I | -1.1% | NA | NA |
| 8 (Control) | A | 0.61 | 10.28 | 0.59 | 0.63 | G-K | NA | -0.3% | NA |
| | N | 0.63 | 9.91 | 0.61 | 0.64 | I-N | 2.7% | **4.3%** | NA |
| | N+P | 0.63 | 10.06 | 0.61 | 0.64 | I-N | 2.5% | 3.3% | NA |
| 14 (Control) | A | 0.62 | 10.28 | 0.60 | 0.63 | H-N | NA | 1.1% | NA |
| | N | 0.62 | 9.91 | 0.60 | 0.63 | H-M | -0.5% | 2.4% | NA |
| | N+P | 0.63 | 10.06 | 0.61 | 0.64 | J-N | 1.5% | 3.7% | NA |
| 21 (Control) | A | 0.61 | 10.28 | 0.59 | 0.62 | G-J | NA | -1.0% | NA |
| | N | 0.64 | 9.91 | 0.62 | 0.65 | N | **5.5%** | **6.3%** | NA |
| | N+P | 0.63 | 10.06 | 0.61 | 0.64 | J-N | 3.4% | **3.5%** | NA |
| 28 (Control) | A | 0.60 | 10.28 | 0.58 | 0.62 | F-H | NA | -1.9% | NA |
| | N | 0.63 | 9.91 | 0.62 | 0.65 | K-N | **5.1%** | **5.0%** | NA |
| | N+P | 0.64 | 10.06 | 0.62 | 0.65 | M-N | **6.0%** | **5.2%** | NA |
| 49 (Control) | A | 0.58 | 10.28 | 0.56 | 0.59 | 0-E | NA | **-5.4%** | NA |
| | N | 0.63 | 9.91 | 0.62 | 0.65 | M-N | **9.7%** | **5.7%** | NA |
| | N+P | 0.63 | 10.06 | 0.62 | 0.65 | L-N | **9.7%** | **4.9%** | NA |
| 65 (Control) | A | 0.57 | 10.28 | 0.55 | 0.58 | 8-B | NA | **-7.2%** | NA |
| | N | 0.60 | 9.91 | 0.58 | 0.61 | E-H | **5.4%** | -0.3% | NA |
| | N+P | 0.62 | 10.21 | 0.60 | 0.63 | H-N | **8.7%** | 2.1% | NA |
| 71 (Control) | A | 0.57 | 10.28 | 0.55 | 0.58 | 9-C | NA | **-7.0%** | NA |
| | N | 0.61 | 10.37 | 0.60 | 0.63 | G-K | **7.3%** | 1.7% | NA |
| | N+P | 0.59 | 10.21 | 0.57 | 0.61 | B-G | 3.8% | -2.4% | NA |
| 76 (Control) | A | 0.56 | 10.28 | 0.54 | 0.58 | 8-10 | NA | **-8.5%** | NA |
| | N | 0.60 | 10.54 | 0.58 | 0.62 | E-H | **7.1%** | -0.1% | NA |
| | N+P | 0.59 | 10.21 | 0.58 | 0.61 | C-G | **5.7%** | -2.1% | NA |

**Table S9 (continuation):** Estimated photochemical efficiency ($F_v/F_m$) for *A. cervicornis* corals exposed to different nutrient treatments and subsequent heat stress. Pairwise comparisons between groups were obtained using Tukey's HSD test (α = 0.05). Stars in *N and *N+P denote the group of corals that were assigned to these treatments, but that were not exposed to elevated nutrients at the time of the measurement. Percentages of change in bold represent comparison among values that were significantly different based on the Tukey's HSD test.

| **Days in the experiment (Phase)** | **Nutrient Treatment** | **$F_v/F_m$ Em mean** | **df** | **Lower CL** | **Upper CL** | **Tukey Group** | **% change respect ambient (same day)** | **% change respect baseline (Day 1)** | **% change respect control (Day 76)** |
|---|---|---|---|---|---|---|---|---|---|
| 84 (Ramp-up) | A | 0.57 | 11.59 | 0.55 | 0.58 | 8-A | NA | **-7.5%** | 1.0% |
| | N | 0.59 | 12.65 | 0.57 | 0.61 | A-G | 4.3% | -1.7% | -1.6% |
| | N+P | 0.60 | 12.02 | 0.58 | 0.62 | D-H | **6.1%** | -0.8% | 1.4% |
| 89 (Ramp-up) | A | 0.55 | 11.59 | 0.53 | 0.56 | 6-9 | NA | **-10.5%** | -2.2% |
| | N | 0.57 | 12.98 | 0.56 | 0.59 | 10-D | **5.0%** | **-4.3%** | -4.2% |
| | N+P | 0.57 | 12.02 | 0.56 | 0.59 | 9-C | 4.4% | **-5.6%** | **-3.5%** |
| 92 (Heat) | A | 0.56 | 11.59 | 0.54 | 0.57 | 7-10 | NA | **-8.7%** | -0.3% |
| | *N | 0.54 | 14.72 | 0.53 | 0.56 | 6-8 | -3.0% | **-9.8%** | -9.7% |
| | *N+P | 0.58 | 12.95 | 0.56 | 0.59 | 0-F | 3.2% | **-4.8%** | -2.7% |
| 96 (Heat) | A | 0.56 | 11.59 | 0.54 | 0.57 | 7-0 | NA | **-8.7%** | -0.2% |
| | *N | 0.50 | 17.30 | 0.48 | 0.52 | 4 | **-10.6%** | **-16.8%** | **-16.7%** |
| | *N+P | 0.53 | 17.25 | 0.51 | 0.55 | 5-7 | -4.8% | **-12.1%** | **-10.2%** |
| 99 (Heat) | A | 0.53 | 11.59 | 0.52 | 0.55 | 5-6 | NA | **-12.9%** | **-4.8%** |
| | *N | 0.41 | 19.16 | 0.39 | 0.42 | 3 | **-23.8%** | **-32.4%** | **-32.3%** |
| | *N+P | 0.48 | 19.12 | 0.47 | 0.50 | 4 | **-9.1%** | **-19.9%** | **-18.2%** |
| 103 (Heat) | A | 0.51 | 11.59 | 0.50 | 0.53 | 4-5 | NA | **-16.0%** | **-8.3%** |
| | *N | 0.39 | 23.21 | 0.37 | 0.41 | 3 | **-23.8%** | **-34.9%** | **-34.8%** |
| | *N+P | 0.41 | 20.27 | 0.39 | 0.43 | 3 | **-20.4%** | **-32.5%** | **-31.0%** |
| 106 (Heat) | A | 0.50 | 11.59 | 0.48 | 0.51 | 4 | NA | **-18.8%** | **-11.3%** |
| | *N | 0.30 | 38.20 | 0.28 | 0.32 | 1 | **-38.7%** | **-49.3%** | **-49.2%** |
| | *N+P | 0.35 | 27.27 | 0.33 | 0.37 | 2 | **-30.1%** | **-42.6%** | **-41.3%** |
| 110 (Heat) | A | 0.40 | 11.59 | 0.38 | 0.42 | 3 | NA | **-34.7%** | **-28.6%** |
| | *N | ND | | | | | NA | NA | NA |
| | *N+P | 0.26 | 83.68 | 0.24 | 0.29 | 1 | **-34.3%** | **-56.6%** | **-55.7%** |

**Table S10:** Estimated photochemical efficiency ($F_v/F_m$) for *O. faveolata* exposed to different nutrient treatments and subsequent heat stress. Pairwise comparisons between groups were obtained using Tukey's HSD test (α = 0.05). Pairwise comparisons between groups were obtained using Tukey's HSD test (α = 0.05). Stars in *N and *N+P denote the group of corals that were assigned to these treatments, but that were not exposed to elevated nutrients at the time of the measurement. Percentages of change in bold represent comparison among values that were significantly different based on the Tukey's HSD test.

| Days in the experiment (Phase) | Nutrient Treatment | $F_v/F_m$ Em mean | df | Lower CL | Upper CL | Tukey Group | % change respect ambient (same day) | % change respect baseline (Day 1) |
|---|---|---|---|---|---|---|---|---|
| 1 (Baseline) | A | 0.51 | 4.29 | 0.47 | 0.54 | H-P | NA | NA |
| | *N | 0.51 | 4.29 | 0.48 | 0.55 | M-Q | 1.6% | NA |
| | *N+P | 0.52 | 4.29 | 0.48 | 0.55 | M-Q | 2.3% | NA |
| 8 (Control) | A | 0.49 | 4.29 | 0.46 | 0.53 | C-N | NA | -2.7% |
| | N | 0.50 | 4.29 | 0.47 | 0.54 | I-N | 2.4% | -2.0% |
| | N+P | 0.50 | 4.29 | 0.46 | 0.54 | D-N | 1.5% | -3.5% |
| 14 (Control) | A | 0.54 | 4.29 | 0.51 | 0.58 | QR | NA | 7.3% |
| | N | 0.54 | 4.29 | 0.50 | 0.57 | O-R | -1.5% | 4.0% |
| | N+P | 0.55 | 4.29 | 0.51 | 0.59 | R | 1.1% | 6.1% |
| 21 (Control) | A | 0.48 | 4.29 | 0.44 | 0.51 | A-I | NA | -5.8% |
| | N | 0.49 | 4.29 | 0.45 | 0.53 | C-M | 2.5% | -5.0% |
| | N+P | 0.49 | 4.29 | 0.46 | 0.53 | C-N | 3.7% | -4.5% |
| 28 (Control) | A | 0.49 | 4.29 | 0.46 | 0.53 | C-N | NA | -2.7% |
| | N | 0.52 | 4.29 | 0.48 | 0.56 | N-R | 5.5% | 1.0% |
| | N+P | 0.54 | 4.29 | 0.50 | 0.57 | P-R | 9.1% | 3.8% |
| 49 (Control) | A | 0.45 | 4.29 | 0.42 | 0.49 | 8-B | NA | -10.4% |
| | N | 0.49 | 4.29 | 0.46 | 0.53 | C-N | 8.7% | -4.2% |
| | N+P | 0.50 | 4.29 | 0.47 | 0.54 | G-O | 11.1% | -2.7% |
| 65 (Control) | A | 0.50 | 4.29 | 0.46 | 0.54 | E-N | NA | -1.1% |
| | N | 0.48 | 4.29 | 0.44 | 0.52 | B-L | -4.1% | -6.6% |
| | N+P | 0.51 | 4.29 | 0.48 | 0.55 | K-Q | 2.6% | -0.8% |
| 71 (Control) | A | 0.47 | 4.29 | 0.43 | 0.50 | 0-D | NA | -7.6% |
| | N | 0.47 | 4.29 | 0.44 | 0.51 | A-H | 1.5% | -7.7% |
| | N+P | 0.45 | 4.29 | 0.41 | 0.48 | 7-A | -4.6% | -13.9% |
| 76 (Control) | A | 0.47 | 4.29 | 0.43 | 0.50 | 0-D | NA | -7.7% |
| | N | 0.47 | 4.29 | 0.44 | 0.51 | A-G | 1.2% | -8.1% |
| | N+P | 0.47 | 4.29 | 0.43 | 0.51 | 9-E | 0.5% | -9.4% |

**Table S10 (continuation):** Estimated photochemical efficiency ($F_v/F_m$) for *O. faveolata* exposed to different nutrient treatments and subsequent heat stress. Pairwise comparisons between groups were obtained using Tukey's HSD test (α = 0.05). Pairwise comparisons between groups were obtained using Tukey's HSD test (α = 0.05). Stars in *N and *N+P denote the group of corals that were assigned to these treatments, but that were not exposed to elevated nutrients at the time of the measurement. Percentages of change in bold represent comparison among values that were significantly different based on the Tukey's HSD test.

| Days in the experiment (Phase) | Nutrient Treatment | $F_v/F_m$ Em mean | df | Lower CL | Upper CL | Tukey Group | % change respect ambient (A) (same day) | % change respect baseline (day 1) | % change respect control (day 76) |
|---|---|---|---|---|---|---|---|---|---|
| 84 (Ramp-up) | A | 0.49 | 4.95 | 0.46 | 0.53 | C-N | NA | -2.8% | 5.4% |
| | N | 0.49 | 4.95 | 0.46 | 0.53 | C-N | 0.2% | -4.2% | 4.3% |
| | N+P | 0.51 | 4.95 | 0.47 | 0.54 | G-P | 2.7% | -2.4% | 7.7% |
| 89 (Ramp-up) | A | 0.48 | 4.95 | 0.44 | 0.51 | 10-K | NA | -5.6% | 2.2% |
| | N | 0.46 | 5.10 | 0.43 | 0.50 | 8-F | -2.8% | -9.7% | -1.7% |
| | N+P | 0.49 | 4.95 | 0.45 | 0.53 | C-N | 2.6% | -5.3% | 4.4% |
| 92 (Heat) | A | 0.52 | 5.10 | 0.48 | 0.55 | L-R | NA | 1.9% | 10.4% |
| | *N | 0.50 | 5.25 | 0.46 | 0.53 | C-N | -3.6% | -3.3% | 5.2% |
| | *N+P | 0.50 | 5.10 | 0.46 | 0.53 | C-N | -3.8% | -4.2% | 5.7% |
| 96 (Heat) | A | 0.43 | 5.10 | 0.40 | 0.47 | 6-9 | NA | -14.3% | -7.1% |
| | *N | 0.44 | 5.25 | 0.40 | 0.47 | 7-0 | 0.9% | -14.9% | -7.4% |
| | *N+P | 0.45 | 5.30 | 0.41 | 0.49 | 7-B | 3.8% | -13.0% | -4.0% |
| 99 (Heat) | A | 0.43 | 5.10 | 0.40 | 0.47 | 5-9 | NA | -14.4% | -7.2% |
| | *N | 0.41 | 5.25 | 0.37 | 0.45 | 3-7 | -5.5% | -20.4% | -13.4% |
| | *N+P | 0.44 | 5.94 | 0.40 | 0.47 | 6-A | 1.1% | -15.4% | -6.7% |
| 103 (Heat) | A | 0.41 | 5.10 | 0.37 | 0.45 | 3-7 | NA | -18.8% | -12.1% |
| | *N | 0.40 | 5.25 | 0.36 | 0.43 | 2-6 | -3.2% | -22.7% | -15.9% |
| | *N+P | 0.43 | 5.94 | 0.39 | 0.46 | 4-8 | 3.6% | -17.8% | -9.3% |
| 106 (Heat) | A | 0.38 | 5.25 | 0.34 | 0.41 | 1-3 | NA | -25.3% | -19.0% |
| | *N | 0.35 | 5.45 | 0.31 | 0.39 | 1 | -7.6% | -32.0% | -26.0% |
| | *N+P | 0.36 | 5.94 | 0.33 | 0.40 | 1-2 | -3.8% | -29.8% | -22.5% |
| 110 (Heat) | A | 0.41 | 5.64 | 0.38 | 0.45 | 3-7 | NA | -18.1% | -11.2% |
| | *N | 0.35 | 5.65 | 0.32 | 0.39 | 1 | **-15.2%** | **-31.7%** | **-25.6%** |
| | *N+P | 0.39 | 5.94 | 0.35 | 0.43 | 1-5 | -5.9% | -24.7% | -16.9% |
| 113 (Heat) | A | 0.42 | 6.23 | 0.39 | 0.46 | 4-8 | NA | -16.3% | -9.3% |
| | N | 0.37 | 5.94 | 0.33 | 0.40 | 1-2 | **-13.8%** | **-29.0%** | **-22.7%** |
| | N+P | 0.39 | 5.94 | 0.35 | 0.42 | 1-4 | -8.7% | -25.3% | -17.6% |

**Table S11:** Estimated photochemical efficiency ($F_v/F_m$) for *S. siderea* corals exposed to different nutrient treatments and subsequent heat stress. Pairwise comparisons between groups were obtained using Tukey's HSD test (α = 0.05). Stars in *N and *N+P denote the group of corals that were assigned to these treatments, but that were not exposed to elevated nutrients at the time of the measurement. Percentages of change in bold represent comparison among values that were significantly different based on the Tukey's HSD test.

| Days in the experiment (Phase) | Nutrient Treatment | $F_v/F_m$ Em mean | df | Lower CL | Upper CL | Tukey Group | % change respect ambient (A) (same day) | % change respect baseline (Day 1) |
|---|---|---|---|---|---|---|---|---|
| 1 (Baseline) | A | 0.47 | 22.59 | 0.45 | 0.49 | B-H | NA | NA |
| | * N | 0.44 | 22.18 | 0.42 | 0.46 | 9-C | -5.4% | NA |
| | * N+P | 0.46 | 23.03 | 0.44 | 0.48 | A-G | -1.6% | NA |
| 8 (Control) | A | 0.49 | 22.59 | 0.48 | 0.51 | G-L | NA | 5.9% |
| | N | 0.50 | 22.18 | 0.48 | 0.52 | H-L | 0.7% | 12.8% |
| | N+P | 0.53 | 23.03 | 0.51 | 0.55 | K-M | 6.5% | 14.6% |
| 14 (Control) | A | 0.55 | 22.59 | 0.53 | 0.57 | M | NA | **18.0%** |
| | N | 0.55 | 22.18 | 0.53 | 0.57 | M | -0.7% | **23.9%** |
| | N+P | 0.56 | 23.03 | 0.54 | 0.57 | M | 0.8% | **20.9%** |
| 21 (Control) | A | 0.49 | 22.59 | 0.47 | 0.51 | F-K | NA | 4.9% |
| | N | 0.51 | 22.18 | 0.49 | 0.52 | I-L | 3.3% | **14.5%** |
| | N+P | 0.52 | 23.03 | 0.50 | 0.54 | J-M | 7.0% | **14.0%** |
| 28 (Control) | A | 0.53 | 22.59 | 0.51 | 0.55 | L-M | NA | **13.6%** |
| | N | 0.55 | 22.18 | 0.53 | 0.57 | M | 3.9% | **24.9%** |
| | N+P | 0.55 | 23.03 | 0.53 | 0.57 | M | 3.5% | **19.5%** |
| 49 (Control) | A | 0.49 | 22.59 | 0.47 | 0.51 | E-J | NA | 4.4% |
| | N | 0.49 | 22.18 | 0.47 | 0.51 | G-J | 0.3% | **10.8%** |
| | N+P | 0.49 | 23.03 | 0.47 | 0.51 | E-I | -0.3% | 5.8% |
| 65 (Control) | A | 0.49 | 23.01 | 0.47 | 0.51 | G-K | NA | 5.4% |
| | N | 0.45 | 22.18 | 0.43 | 0.47 | 0-D | -9.3% | 1.0% |
| | N+P | 0.48 | 23.46 | 0.46 | 0.50 | D-I | -2.0% | 5.0% |
| 71 (Control) | A | 0.48 | 23.01 | 0.46 | 0.50 | D-I | NA | 3.1% |
| | N | 0.46 | 22.18 | 0.44 | 0.48 | A-G | -4.4% | 4.3% |
| | N+P | 0.47 | 23.03 | 0.46 | 0.49 | C-I | -1.4% | 3.3% |
| 76 (Control) | A | 0.48 | 23.01 | 0.46 | 0.50 | D-I | NA | 3.3% |
| | N | 0.46 | 22.18 | 0.44 | 0.48 | A-H | -4.1% | 4.8% |
| | N+P | 0.47 | 23.03 | 0.45 | 0.49 | C-I | -2.0% | 2.9% |

**Table S11 (continuation):** Estimated photochemical efficiency ($F_v /F_m$) in *S. siderea* corals exposed to different nutrient treatments and subsequent heat stress. Pairwise comparisons between groups were obtained using Tukey's HSD test (α = 0.05). Stars in *N and *N+P denote the group of corals that were assigned to these treatments, but that were not exposed to elevated nutrients at the time of the measurement. Percentages of change in bold represent comparison among values that were significantly different based on the Tukey's HSD test.

| Days in the experiment (Phase) | Nutrient Treatment | $F_v/F_m$ Em mean | df | Lower CL | Upper CL | Tukey Group | % change respect ambient (A) (same day) | % change respect Baseline (Day 1) | % change respect Control (day 76) |
|---|---|---|---|---|---|---|---|---|---|
| 84 (Ramp- up) | A | 0.47 | 31.95 | 0.45 | 0.49 | B-I | NA | 0.5% | -2.7% |
| | N | 0.43 | 30.17 | 0.41 | 0.45 | 8-B | -7.8% | -2.1% | -6.6% |
| | N+P | 0.47 | 31.05 | 0.45 | 0.49 | B-I | 0.0% | 2.1% | -0.7% |
| 89 (Ramp- up) | A | 0.44 | 31.95 | 0.42 | 0.46 | 9-C | NA | -5.8% | -8.9% |
| | N | 0.43 | 30.17 | 0.41 | 0.45 | 7-A | -3.2% | -3.6% | -8.0% |
| | N+P | 0.42 | 31.05 | 0.40 | 0.44 | 6-A | -3.6% | -7.8% | -10.4% |
| 92 (Heat) | A | 0.45 | 31.95 | 0.43 | 0.47 | 9-E | NA | -3.9% | -7.0% |
| | * N | 0.45 | 30.17 | 0.43 | 0.47 | 0-F | 0.3% | 1.9% | -2.8% |
| | * N+P | 0.45 | 31.05 | 0.43 | 0.47 | 9-E | -0.3% | -2.7% | -5.4% |
| 96 (Heat) | A | 0.38 | 31.95 | 0.36 | 0.40 | 1-5 | NA | -19.5% | **-22.1%** |
| | * N | 0.42 | 30.17 | 0.40 | 0.44 | 5-0 | 11.2% | -5.4% | **-9.7%** |
| | * N+P | 0.39 | 31.05 | 0.37 | 0.41 | 2-8 | 4.6% | -14.5% | **-16.8%** |
| 99 (Heat) | A | 0.38 | 31.95 | 0.35 | 0.40 | 1-5 | NA | -19.6% | **-22.1%** |
| | * N | 0.41 | 30.17 | 0.39 | 0.43 | 3-0 | 9.1% | -7.2% | **-11.4%** |
| | * N+P | 0.38 | 31.05 | 0.36 | 0.40 | 1-5 | 0.8% | -17.6% | **-19.9%** |
| 103 (Heat) | A | 0.35 | 31.95 | 0.33 | 0.37 | 1-2 | NA | -24.3% | **-26.7%** |
| | * N | 0.41 | 30.17 | 0.39 | 0.43 | 3-9 | **15.0%** | -8.0% | **-12.2%** |
| | * N+P | 0.37 | 31.05 | 0.35 | 0.39 | 1-3 | 4.1% | -20.0% | **-22.2%** |
| 106 (Heat) | A | 0.34 | 31.95 | 0.32 | 0.36 | 1 | NA | -26.7% | **-29.0%** |
| | * N | 0.37 | 30.17 | 0.35 | 0.39 | 1-4 | 8.7% | -15.8% | **-19.6%** |
| | * N+P | 0.34 | 31.05 | 0.32 | 0.36 | 1 | 0.0% | -25.5% | **-27.6%** |
| 110 (Heat) | A | 0.37 | 31.95 | 0.35 | 0.39 | 1-4 | NA | -20.5% | **-23.0%** |
| | * N | 0.41 | 30.17 | 0.39 | 0.43 | 4-10 | 10.7% | -6.9% | **-11.2%** |
| | * N+P | 0.38 | 31.05 | 0.36 | 0.40 | 1-6 | 3.0% | -16.8% | **-19.1%** |
| 113 (Heat) | A | 0.37 | 31.95 | 0.35 | 0.39 | 1-4 | NA | -20.1% | **-22.7%** |
| | * N | 0.42 | 30.17 | 0.40 | 0.44 | 5-10 | **12.2%** | -5.3% | **-9.6%** |
| | * N+P | 0.38 | 31.05 | 0.36 | 0.40 | 1-7 | 2.7% | -16.6% | **-19.0%** |

**Table S12:** Estimated chlorophyll-*a* content (µg $cm^{-2}$) before and after heat stress in *A. cervicornis*, *O. faveolata* and *S. siderea* exposed to different nutrient treatments. Pairwise comparisons between groups were obtained using Tukey's HSD test (α = 0.05). Stars in *N and *N+P denote the group of corals that were assigned to these treatments, but that were not exposed to elevated nutrients at the time of the measurement. Percentages of change in bold represent comparison among values that were significantly different based on the Tukey's HSD test. Each coral species was evaluated separately (Tukey groups compare values inside each species, but not across species).

| Coral species | Days in the experiment | Nutrient Treatment | Estimated Mean | SE | n | Tukey Group | % change respect ambient (A) (same day) | % change respect control (day 78) |
|---|---|---|---|---|---|---|---|---|
| *A.cer* | 78 (Control) | A | 3.58 | 0.4 | 7 | 2 | NA | NA |
| | | N | 5.12 | 0.37 | 8 | 3 | **43.1%** | NA |
| | | N+P | 4.97 | 0.40 | 8 | 3 | **38.9%** | NA |
| | 113 (Heat) | A | 1.35 | 0.37 | 8 | 1 | NA | **-62.2%** |
| | | * N | ND | NA | 0 | NA | NA | NA |
| | | * N+P | 0.42 | 1.05 | 1 | 1-2 | -68.9% | **-91.5%** |
| *O.fav* | 78 (Control) | A | 3.82 | 0.63 | 8 | 2 | NA | NA |
| | | N | 5.69 | 0.63 | 8 | 3 | **49.2%** | NA |
| | | N+P | 5.90 | 0.63 | 8 | 3 | **54.6%** | NA |
| | 113 (Heat) | A | 2.29 | 0.68 | 6 | 12 | NA | -40.0% |
| | | * N | 1.52 | 0.68 | 6 | 1 | -33.5% | **-73.2%** |
| | | * N+P | 2.77 | 0.71 | 5 | 12 | 20.8% | **-53.1%** |
| *S.sid* | 78 (Control) | A | 4.49 | 0.45 | 14 | 3 | NA | NA |
| | | N | 5.37 | 0.46 | 14 | 4 | **19.77%** | NA |
| | | N+P | 5.36 | 0.46 | 13 | 4 | **19.6%** | NA |
| | 113 (Heat) | A | 0.81 | 0.46 | 14 | 1 | NA | **-81.9%** |
| | | * N | 1.69 | 0.47 | 14 | 2 | **108.6%** | **-68.4%** |
| | | * N+P | 1.69 | 0.46 | 14 | 2 | **108.2%** | **-68.5%** |

**Table S13:** Estimated Symbiodiniaceae areal density (Symbiodiniaceae cells $cm^{-2}$) before and after heat stress in *A. cervicornis*, *O. faveolata* and *S. siderea* exposed to different nutrient treatments. Pairwise comparisons between groups were obtained using Tukey's HSD test (α = 0.05). Stars in *N and *N+P denote the group of corals that were assigned to these treatments, but that were not exposed to elevated nutrients at the time of the measurement. Percentages of change in bold represent comparison among values that were significantly different based on the Tukey's HSD test. Each coral species was evaluated in a separated model (Tukey groups compare values inside each species, but not across species). ND: No data collected due coral mortality.

| Coral species | Days in the experiment (Temperature) | Nutrient Treatment | Estimated Mean | SE | n | Tukey Group | % change respect ambient (A) (same day) | % change respect control (day 78) |
|---|---|---|---|---|---|---|---|---|
| *A.cer* | 78 (Control) | A | 4.1 | 0.5 | 7 | 2 | NA | NA |
| | | N | 4.8 | 0.5 | 8 | 2 | 18.1% | NA |
| | | N+P | 5.3 | 0.5 | 8 | 2 | 29.7% | NA |
| | 113 (Heat) | A | 1.1 | 0.5 | 8 | 1 | NA | **-73.9%** |
| | | * N | ND | NA | 0 | NA | NA | NA |
| | | * N+P | 0.3 | 1.01 | 1 | 1 | -73.5% | **-94.7%** |
| *O.fav* | 78 (Control) | A | 1.56 | 0.17 | 8 | 2 | NA | NA |
| | | N | 1.97 | 0.17 | 8 | 2 | 26.3% | NA |
| | | N+P | 2.01 | 0.17 | 8 | 2 | 28.8% | NA |
| | 113 (Heat) | A | 0.77 | 0.20 | 6 | 1 | NA | **-50.6%** |
| | | * N | 0.59 | 0.20 | 6 | 1 | -23.4% | **-70.1%** |
| | | * N+P | 0.82 | 0.21 | 5 | 1 | 6.5% | **-59.2%** |
| *S.sid* | 78 (Control) | A | 1.20 | 0.09 | 14 | 2 | NA | NA |
| | | N | 1.42 | 0.09 | 14 | 2 | 17.8% | NA |
| | | N+P | 1.40 | 0.10 | 13 | 2 | 16.5% | NA |
| | 113 (Heat) | A | 0.36 | 0.09 | 14 | 1 | NA | **-70.3%** |
| | | * N | 0.54 | 0.09 | 14 | 1 | 52.5% | **-61.6%** |
| | | * N+P | 0.42 | 0.09 | 14 | 1 | 19.1% | **-69.7%** |

**Table S14:** Estimated Chlorophyll-*a* concentration and Symbiodiniaceae areal density (Symbiodiniaceae cells $cm^{-2}$) before and after heat stress in *S. siderea*. Pairwise comparisons between groups were obtained using Tukey's HSD test (α = 0.05). Percentages of change in bold represent comparison among values that were significantly different based on the Tukey's HSD test.

| Days in the experiment (Temperature) | Dominant symbiont | Estimated Mean | SE | Tukey Group | % change respect control (day 78) |
|---|---|---|---|---|---|
| | | Chlorophyll-*a* | | | |
| 78 (Control) | *Cladocopium* C3 | 6.78 | 0.66 | 4 | NA |
| | *Cladocopium* C1 | 3.93 | 0.63 | 2-3 | NA |
| | *D. trenchii* | 4.38 | 0.49 | 3 | NA |
| 113 (Heat) | *Cladocopium* C3 | 1.61 | 0.69 | 1-2 | **-73.3%** |
| | *Cladocopium* C1 | 1.04 | 0.62 | 1 | **-73.6%** |
| | *D. trenchii* | 1.51 | 0.47 | 1 | **-65.5%** |
| | | Sybiodiniaceae areal density (cells $cm^{-2}$) | | | |
| 78 (Control) | *Cladocopium* C3 | 1.34 | 0.07 | 4 | NA |
| | *Cladocopium* C1 | 1.07 | 0.07 | 3-4 | NA |
| | *D. trenchii* | 1.06 | 0.05 | 3 | NA |
| 113 (Heat) | *Cladocopium* C3 | 0.43 | 0.07 | 1-2 | **-68.0%** |
| | *Cladocopium* C1 | 0.60 | 0.07 | 1 | **-44.1%** |
| | *D. trenchii* | 0.72 | 0.05 | 1 | **-32.4%** |
